# RAF isoform selectivity of MEK inhibitors and rational design of a covalent ARAF-MEK inhibitor

**DOI:** 10.64898/2025.12.15.693503

**Authors:** Sayan Chakraborty, Jie Jiang, Javier Vinals Camallonga, Dimitris Gazgalis, Thomas W. Gero, Maurício T. Tavares, Georgia Dittemore, Emre Tkacik, Dong Man Jang, Byung Hak Ha, Felix H. Gottlieb, Talya Levitz, Rebecca J. Metivier, Anna M. Schmoker, David A. Scott, Hyesung Jeon, Pasi A. Jänne, Michael J. Eck

## Abstract

Aberrant activation of the RAS/RAF/MEK/ERK pathway is a frequent cause of cancer. Allosteric MEK inhibitors block this pathway by binding RAF-MEK complexes to prevent activation of MEK by RAF. However, how MEK inhibitor potency varies across the three RAF isoforms remains poorly understood. We profiled seven allosteric MEK inhibitors and observed a striking hierarchy of sensitivity: all most potently inhibited CRAF-driven MEK activation while relatively sparing ARAF-driven activation. We identified point mutations in ARAF and CRAF proximate to the MEK inhibitor binding site that markedly altered inhibitor sensitivity. Using a rational design approach, we developed a more potent inhibitor of ARAF-driven MEK signaling, TWG-07-148. Our cryo-EM structure shows how this acrylamide-containing analog of MEK inhibitor trametinib covalently targets Cys514, a residue unique to ARAF. Our studies highlight the importance of the activating RAF isoform as a determinant of MEK inhibitor sensitivity and provide proof-of-concept for development of MEK inhibitors that more effectively block ARAF-driven MEK signaling via covalent targeting of Cys514.

## Introduction

MEK is the central kinase in the three-tiered RAF/MEK/ERK (MAP kinase) cascade (*1–3*). This signaling axis is normally triggered in response to activation of receptor tyrosine kinases on the cell surface, leading to activation of RAS, which in turn engages and activates RAF, the kinase at the top of this cascade (*2, 4, 5*). Physiologic activation of the pathway results in regulated changes in cell proliferation, differentiation and survival, but mutations in components of this pathway can lead to its uncontrolled activation and are a frequent cause of cancer (*2, 3, 6, 7*). RAS is the most frequently mutated oncogenic driver (*8–10*). Mutations in RAS are estimated to occur in 19% of all cancers (*10, 11*), and mutations in BRAF, one of three RAF isoforms, drive approximately 7% of all cancers (*12*). By far the most common activating mutation in BRAF is the V600E point mutation (*13*), which accounts for approximately half of all malignant melanoma (*14*) and as much as 60% (*15*) of papillary thyroid carcinoma and is also a less frequent driver of lung, colorectal, and other cancers (*12, 16*).

MEK inhibitors are a valuable therapeutic for treatment of certain tumors arising from aberrant activation of this pathway (*17, 18*). MEK inhibitors trametinib, cobimetinib, and binimetinib are used in combination with RAF inhibitors dabrafenib, vemurafenib, and encorafenib, respectively, to treat BRAF^V600E+^ melanomas (*19–23*). Selected RAF and MEK inhibitor combinations are also approved for treatment of other solid tumors driven by BRAF^V600E^ (*20, 22, 24, 25*). MEK inhibitors selumetinib and mirdametinib are used to treat plexiform neurofibromas in patients with type I neurofibromatosis (*26*), a condition caused by loss of function mutations in NF1, a GTPase-activating protein that down-regulates RAS.

These and other clinical stage MEK inhibitors differ from the vast majority of kinase inhibitors in that they have an allosteric mechanism of action (*27, 28*). Rather than blocking binding of ATP and directly inhibiting the ability of MEK to phosphorylate ERK, they bind in an allosteric pocket where they stabilize a conformation of the MEK activation segment that interferes with the ability of RAFs to phosphorylate and activate MEK (*28*). This allosteric mechanism of action has been recognized almost from the time of the discovery of the first MEK inhibitors in the mid-1990’s (*27*), and its general structural basis became clear with the first co-crystal structures in 2004 (*28*). More recently, structural studies of MEK inhibitors bound to RAF-MEK complexes and with MEK in complex with KSR (a RAF-like pseudokinase) have revealed that the MEK inhibitor binding pocket is formed in part by the juxtaposed RAF kinase, and that MEK inhibitors can differentially stabilize these complexes (*29–32*).

Owing to their allosteric mechanism, the pharmacologic effects of MEK inhibitors can be idiosyncratic. In particular, they can vary in their efficacy depending on how the pathway is activated. For example, cobimetinib and mirdametinib are effective in blocking MAP kinase signaling in the context of BRAF^V600E^, but not mutant KRAS, while MEK inhibitors GDC-0623 and trametinib are effective in both oncogenic contexts (*33, 34*). Activation of CRAF in KRAS mutant cells was identified as a determinant of sensitivity to mirdametinib – MEK was found to be less susceptible to inhibition when activated by CRAF as compared with BRAF^V600E^ (*34*). While MEK inhibitors act most potently on RAF-MEK complexes to block MEK activation, at higher concentrations most can inhibit active, phosphorylated MEK (*29, 35*). This property may explain in part the differing ability of MEK inhibitors to block ERK activation in KRAS mutant cells (*29, 35*). One exception is avutometinib (CH5126766) which is known to selectively target RAF-MEK complexes and to have essentially no ability to inhibit free, active MEK (*29, 36*).

Although MEK inhibitors have long been known to target RAF-MEK complexes, to our knowledge their activity against complexes containing different RAF isoforms has not been systematically investigated. This question has important therapeutic implications. Although rare in comparison with BRAF mutations, activating mutations in both ARAF and CRAF have also been identified as oncogenic drivers (*37–41*). Additionally, RAF isoforms can compensate for one another during targeted therapy and can mediate resistance to BRAF and MEK inhibitors that arises via relief of negative feedback inhibition (*42, 43*). In this study, we systematically profiled seven allosteric MEK inhibitors (MEKi) across RAF isoforms using both biochemical and cell-based assays. We discovered marked differences in isoform-selectivity with most MEK inhibitors exhibiting relative sparing of ARAF-driven MEK signaling. Through structure-guided mutagenesis, we identified RAF residues in the proximity of the MEK inhibitor binding pocket — Gly513 and Cys514 in ARAF and Asn553 in CRAF — as critical determinants of sensitivity to MEK inhibitors. Recognizing the proximity of Cys514 in ARAF to the MEK inhibitor binding site, we explored covalent targeting of this residue leading to the development of TWG-07-148, an acrylamide-containing analog of trametinib. TWG-07-148 binds irreversibly to Cys514 on ARAF, and more potently inhibits ARAF-MEK signaling as compared with trametinib and other existing MEK inhibitors. Collectively, our studies reveal a hierarchy of RAF isoform sensitivity to current MEK inhibitors, with CRAF>BRAF>ARAF, and provide a strategy for more potent inhibition of the ARAF-driven MEK activation via covalent targeting of a cysteine residue that is unique to ARAF.

## Results

### MEK inhibitors differ in potency across RAF isoforms

To test MEK inhibitor potency in an isogenic context, we prepared Ba/F3 cell lines that stably express activated alleles of ARAF, BRAF or CRAF (ARAF^S214A^, BRAF^S365A^, CRAF^S259A^). These mutations ablate the negative-regulatory phosphorylation and 14-3-3 binding site in each RAF and all promoted IL-3 independent proliferation of the respective Ba/F3 cell line. We also prepared Ba/F3 cells transformed by BRAF^V600E^. We tested a panel of 7 MEKi on proliferation of each cell line and on the parental Ba/F3 cells using the CellTiter-Glo (Promega) cell viability assay (Figure 1A, Supplementary Figure 1A,B). Although Ba/F3 cells endogenously express low levels of wild-type (WT) RAFs, all were sensitized to MEK inhibition as compared with the parental Ba/F3 cells, further confirming their dependence on the activated RAF alleles. All 7 of the MEKi tested were most potent on the CRAF-driven Ba/F3 cells and least potent on ARAF-driven cells (Figure 1A, Supplementary Figure 1B, Supplementary Table 1). While the difference in potency on CRAF versus ARAF varied, it was typically on the order of 10- to 100-fold. Among the MEK inhibitors we tested, trametinib was the most uniformly potent across RAF isoforms, with a sub-nanomolar IC_50_ against CRAF^S259A^ and an IC_50_ of ∼29 nM against ARAF^S214A^. The tested MEK inhibitors exhibited intermediate potency on BRAF^S365A^, with measured IC_50_ values ranging from 3.8 nM for trametinib to ∼700 nM for selumetinib. Potencies on BRAF^S365A^ and BRAF^V600E^ were very similar, indicating that the two different mechanisms of activation do not differentially affect MEK inhibition (Figure 1A, Supplementary Figure 1B, Supplementary Table 1).

**Figure 1.**
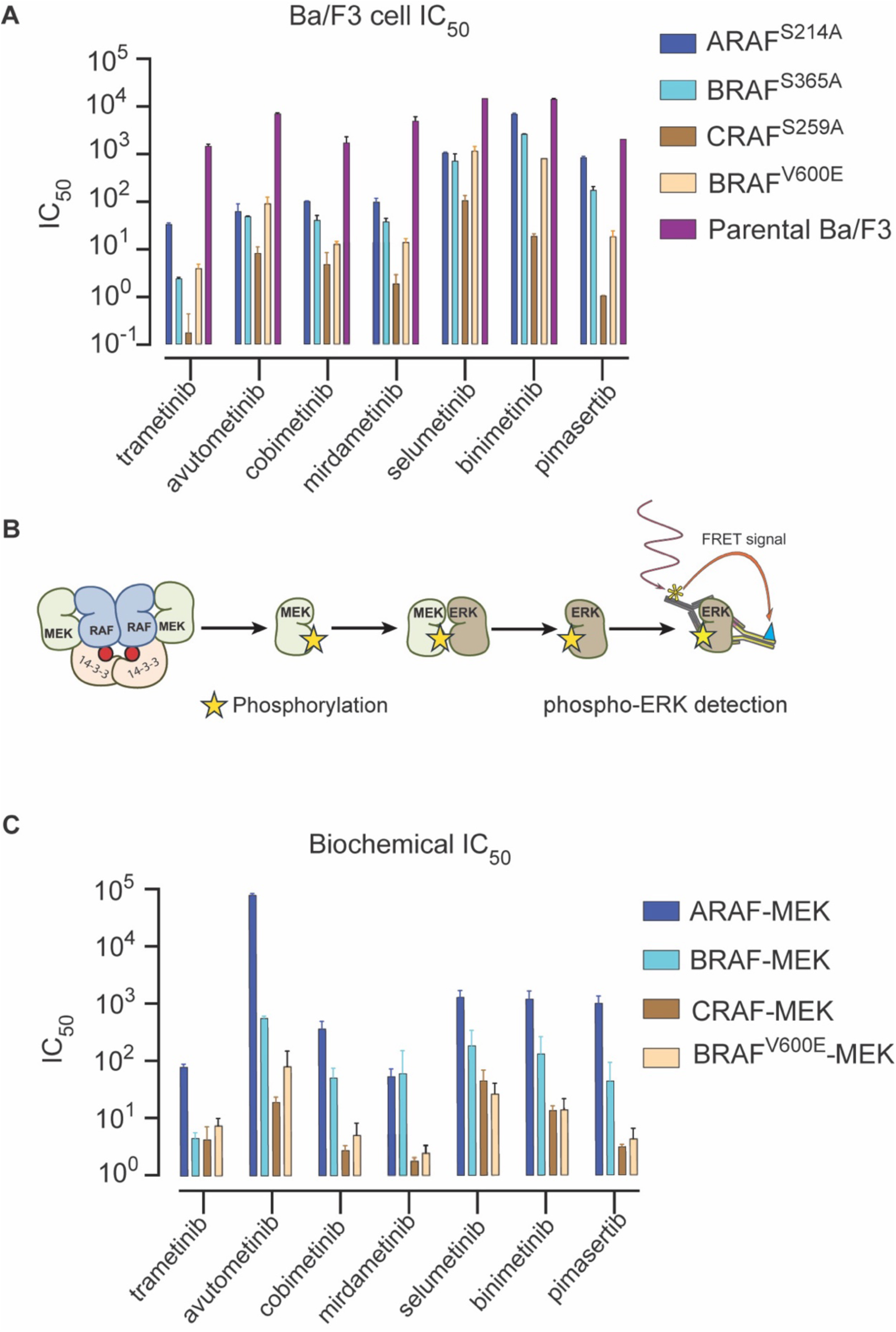
Potency and selectivity of allosteric MEK inhibitors across RAF isoforms. (**A**) Bar chart showing the cellular IC_50_ values of allosteric MEKi in Ba/F3 cells expressing activated alleles of ARAF, BRAF, or CRAF. The parental Ba/F3 cell line lacking exogenous RAF expression was included as a control. IC_50_ values for inhibition of proliferation are reported as mean, N=2 with Standard Error (SE) indicated by error bars. (**B**) Schematic of the cascade biochemical assay used to measure ERK phosphorylation in the presence or absence of MEK inhibitors. (**C**) Bar chart summarizing the biochemical potency of MEKi against purified active RAF-MEK complexes, reported as mean of N=3 with SE indicated by error bars.

We also compared potencies of MEK inhibitors in biochemical assays against purified ARAF-MEK1, BRAF-MEK1 and CRAF-MEK1 complexes. These preparations consist of active, 14-3-3-bound dimers of the respective RAF kinase domains, all prepared as complexes with MEK1 (*44*). For ARAF and CRAF, we incorporated an “SSDD” substitution in their N-terminal acidic motifs (NtA motifs, replacing their native SSYY and SGYY sequences, respectively) which has previously been shown to increase their catalytic activity in lieu of tyrosine phosphorylation of their native NtA motifs(*45*). In this cascade assay, phosphorylation of ERK2 is measured using a TR-FRET-based sandwich immunoassay that employs a phospho-specific antibody for pERK1/2 (T185/Y187) (Figure 1B). Relative potencies in these biochemical assays largely paralleled those in our Ba/F3 assays – all seven MEKi were most potent against the CRAF-MEK complex and least potent against the ARAF-MEK complex (Figure 1C, Supplementary Figure 1C, Supplementary Table 2). As in the cellular assays, differences in potency on ARAF versus CRAF were on the order of 10- to 100-fold. One difference, however, was that most of the tested MEKi were considerably more potent against BRAF^V600E^-MEK than against the wild-type BRAF-MEK complex in the biochemical assay. This difference could arise from our use of a monomeric preparation of BRAF^V600E^ that lacked the 14-3-3-stabilized dimerization of the BRAF^V600E^-MEK complex. Overall, our findings in both the cellular and biochemical assays demonstrate that all 7 MEK inhibitors tested are most potent against CRAF-driven MEK activation and least potent against ARAF-driven MEK activation. While this rank-order potency was the same irrespective of the MEK inhibitor chemotype, the extent of the RAF isoform-selectivity did vary markedly among the inhibitors tested (Supplementary Tables 1, 2).

### Mutations in the RAF αG-helix can confer sensitivity or resistance to MEK inhibition

Crystal structures of RAF-MEK complexes show that the MEK inhibitor binding cleft is bordered on one end by three residues at the N-terminus of the αG-helix in RAF(*29*) (Figure 2A). Both trametinib and avutometinib make direct contacts with Asn^661^ in this segment in BRAF-MEK complexes, and mutations in this region have been shown to affect the ability of trametinib to stabilize RAF-MEK and KSR-MEK complexes(*29, 30*). Sequence comparisons reveal that the Asn-Asn-Arg sequence conserved in BRAF and CRAF is replaced by Gly-Cys-Arg in ARAF (Figure 2B) and given the proximity of these residues to the MEKi binding site and their divergence in ARAF, we were interested to test the effect of mutations in this region on the potency of MEK inhibitors.

**Figure 2.**
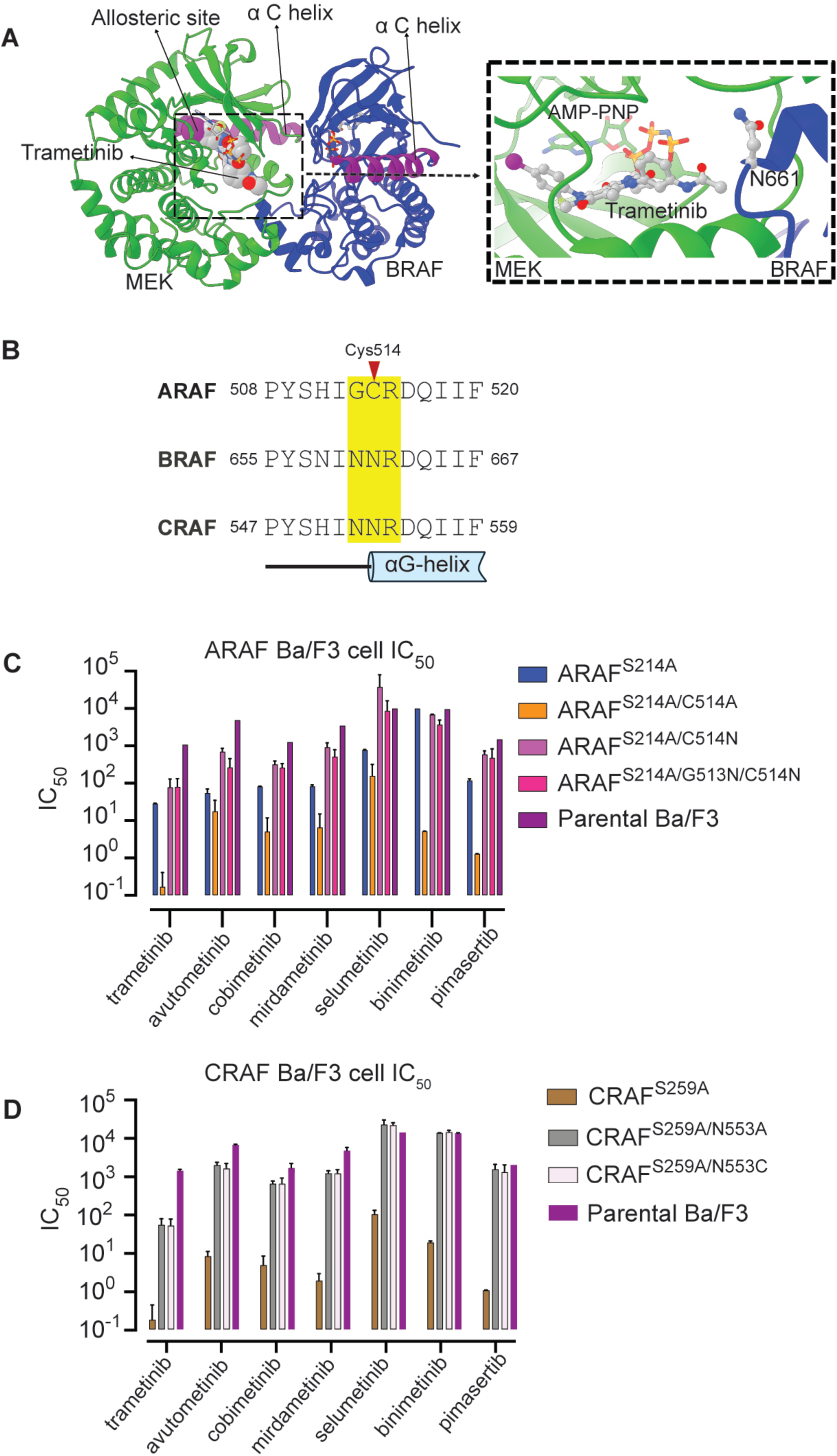
Impact of ARAF and CRAF mutations on MEK inhibitor sensitivity. (**A**) Ribbon diagram depicting the structure of the BRAF-MEK complex, with trametinib bound to the allosteric site of MEK (PDB entry 7M0Y). The inset shows a detailed view of trametinib and its relationship to Asn661 in the BRAF αG helix. (**B**) Aligned sequences of human ARAF, BRAF and CRAF in the region of the RAF-MEK allosteric pocket. Residues that form the RAF portion of the pocket are highlighted in yellow and the position of Cys514 in ARAF is marked with an arrow. (**C** and **D**) Bar charts showing the cellular IC_50_ values of allosteric MEKi in Ba/F3 cells expressing individual ARAF (**C**) or CRAF (**D**) mutants, as indicated. The parental Ba/F3 cell line lacking exogenous RAF expression was included as a control.

We mutated Cys514 in ARAF (corresponding to Asn661 in BRAF) to alanine and asparagine and we also prepared the ARAF G513N/C514N double mutant to match the NNR motif found in BRAF and CRAF. These mutations were introduced into the activated ARAF^S214A^ construct in Ba/F3 cells. Interestingly, the C514A mutation dramatically sensitized ARAF to MEK inhibition. All 7 MEKi were more potent against this mutant, with IC_50_ shifts ranging from ∼250-fold for trametinib, to ∼8-fold for avutometinib (Figure 2C, Supplementary Figure 2A, and Supplementary Table 3). By contrast, the single and double asparagine mutations increased IC_50_’s for the panel of MEK inhibitors (Figure 2C, Supplementary Figure 2B, and Supplementary Table 3). These large shifts in MEKi sensitivity were not recapitulated in our biochemical assay, although the C514A mutant did modestly sensitize to the tested MEK inhibitors (Supplementary Figure 2C, Supplementary Table 4).

To better understand the underlying basis for these changes in inhibitor sensitivity, we carried out molecular dynamics simulations with the WT ARAF-MEK complex and with the C514A and C514N mutants. In all trajectories, we observed coupled movements between the ARAF αG helix and the MEK activation segment helix. With both mutants, we observed closer contact between these structural elements and also between the αG helix and the bound inhibitor (Supplementary Figure 3 A-C). Calculated protein-ligand interaction energies predicted higher affinity for the trametinib scaffold in both mutants as compared with WT (Supplementary Figure 3D). These simulations support the notion that changes in the dynamics of the interface affect inhibitor affinity, and calculated changes in ligand interaction energies are consistent with observed changes in our biochemical, but not cellular, assays.

We also examined the effect of cysteine and alanine mutations in the corresponding residue of CRAF (N553A and N553C) in our Ba/F3 cell model. Both substitutions resulted in profound resistance to MEK inhibition, resulting in sensitivity similar to that of the parental Ba/F3 cells (Figure 2D, Supplementary Figure 2B, Supplementary Table 3). Collectively, these findings reinforce the importance of the RAF partner in RAF-MEK complexes as a determinant of sensitivity to the tested MEK inhibitors.

### A more potent ARAF-MEK inhibitor via covalent targeting of ARAF^C514^

Although ARAF itself is only rarely mutated in cancer (*12, 39, 46*), it is increasingly appreciated as an important therapeutic target owing to its role in mediating resistance to RAF inhibitors, particularly in NRAS-mutant cancers (*40, 47*). As we find above, most MEK inhibitors are least effective in the context of ARAF-driven MEK activation, and furthermore, current RAF inhibitors, particularly type II inhibitors are also least effective against ARAF (*44, 48*). Motivated by this need and encouraged by the position of ARAF Cys514 in the MEK inhibitor binding pocket, we sought to develop a more potent inhibitor of ARAF-MEK via covalent targeting of this residue.

In our rational design approach, we appended covalent warheads (acrylamide or chloroacetamide) to MEK inhibitor scaffolds in a manner expected to juxtapose the warheads with the target cysteine residue in ARAF. We synthesized a series of 11 compounds based on the core scaffolds of trametinib, avutometinib, cobimetinib and selumetinib, and assessed their ability to form the desired covalent bond with ARAF as well as their potency against activated ARAF^S214A^ in the Ba/F3 cell assay (Figure 3, Supplementary Figure 4 and 5). Selected acrylamide and chloroacetamide derivatives of both trametinib and avutometinib stoichiometrically labeled ARAF when applied to the ARAF-MEK complex (Figure 3). Intact mass analysis suggested a single site of labeling at ∼100% occupancy for both trametinib-based TWG-07-148 and avutometinib-based GD-01-012 (Figure 4A) and mapping of the sites by LC-MS/MS analysis of a tryptic digest of the labeled samples confirmed the site of labeling as Cys514 in ARAF, the target residue (Figure 4B). The trametinib-based compounds were potent in the ARAF-driven Ba/F3 assay, with TWG-07-148 exhibiting an IC_50_ = 5.5 nM, approximately 5-fold more potent that trametinib (Figure 4C, Supplementary Table 1). TWG-07-148 was also more potent than trametinib in the biochemical ARAF-MEK assay (Figure 4D).

**Figure 3.**
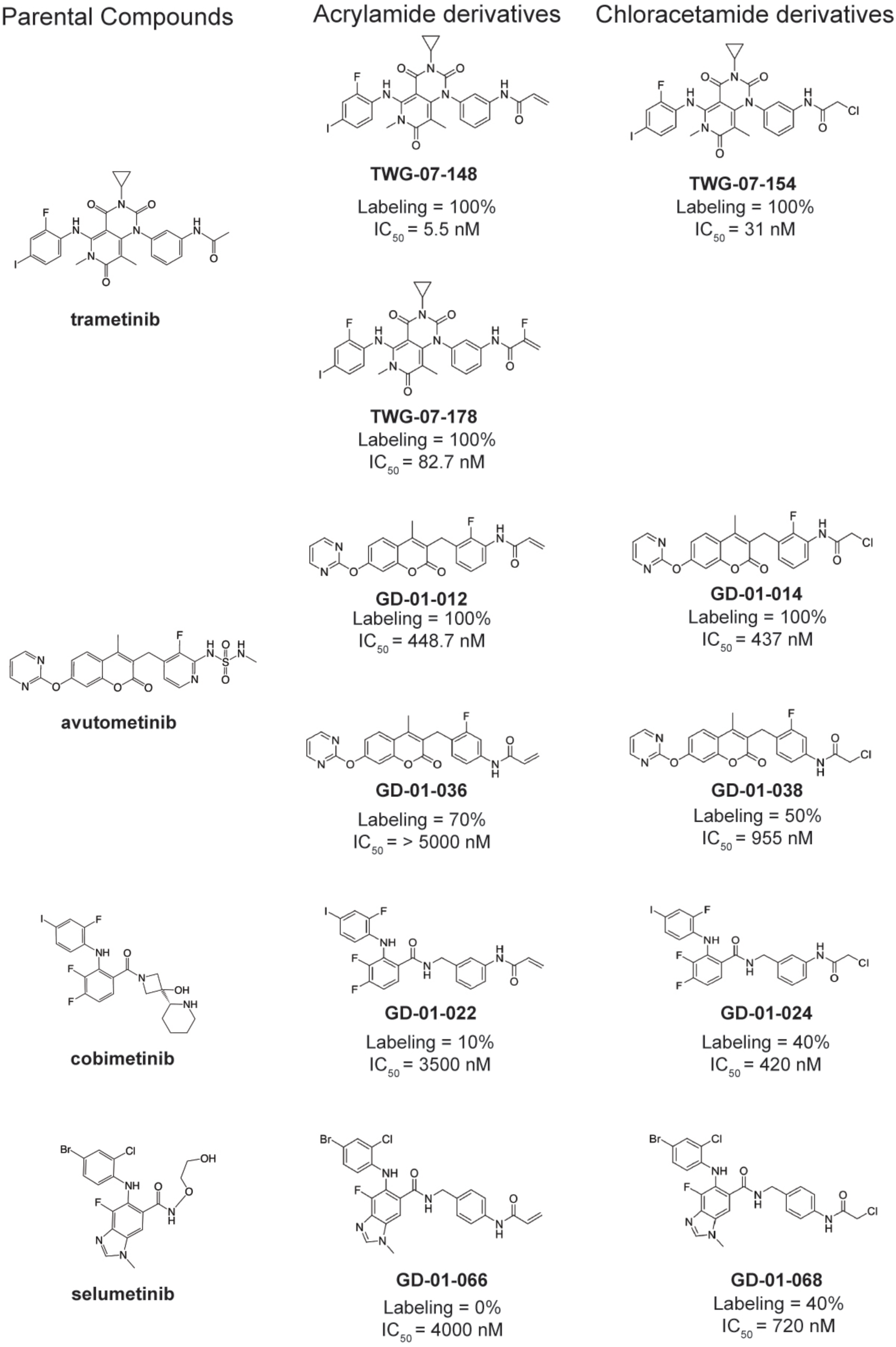
Structures of covalent MEK inhibitors targeting ARAF C514. Electrophilic acrylamide or chloroacetamide groups were introduced in existing allosteric MEK inhibitor scaffolds (those of trametinib, avutometinib, cobimetinib, and selumetinib) with the goal of covalent targeting of Cys514 in ARAF. For each of the resulting compounds, covalent labeling of ARAF was measured in an intact mass spectrometry assay with the ARAF-MEK complex (Labeling %). IC_50_ values provided for each compound are for inhibition of proliferation of Ba/F3 cells expressing ARAF^S214A^.

**Figure 4.**
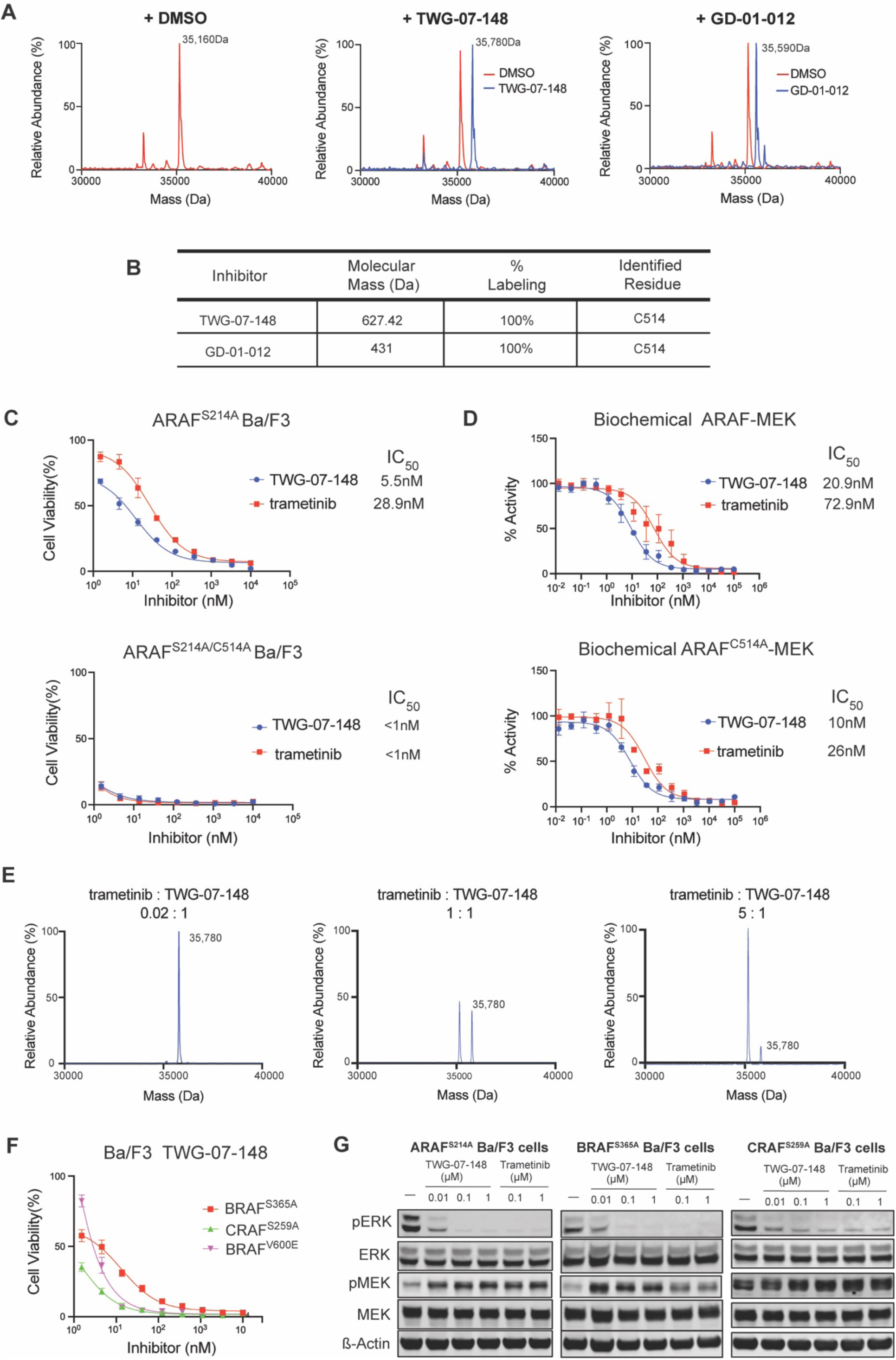
Biochemical and cellular evaluation of covalent ARAF-MEK inhibitor TWG-07-148. (**A**) Intact mass spectrometry analysis of the ARAF-MEK1 complex after treatment with trametinib derivative TWG-07-148 (center panel), avutometinib derivative GD-01-012 (right panel) or DMSO control (left panel). The DMSO control is superimposed in red for reference in the TWG-07-148 and GD-01-012 panels. Note that MEK1 lies outside the mass range illustrated. (**B**) LC/MS/MS analysis of a tryptic digest of the TWG-07-148 and GD-01-012 treated samples identified Cys514 as the site of covalent modification. (**C**) Dose-response curves for TWG-07-148 and trametinib in Ba/F3 cells expressing ARAF^S214A^ (upper panel) or ARAF^S214A/C514A^ (lower panel). (**D**) Dose-response curves for TWG-07-148 and trametinib in the biochemical cascade assay with ARAF-MEK1 (upper panel) or ARAF^C514A^-MEK1 (lower panel). (**E**) Mass spectrometry-based competition assay with trametinib and TWG-07-148. Increasing concentrations of the reversible MEK inhibitor trametinib compete for the same pocket as TWG-07-148 to block labeling of ARAF Cys514, supporting MEK-mediated delivery of the covalent compound to Cys514 of ARAF. The ARAF-MEK1 complex (3 μM) was treated with an equimolar concentration of TWG-07-148 plus trametinib at 0.06 μM (0.02:1), 3 μM (1:1) or 15 μM (5:1) and analyzed by intact mass spectrometry. (**F**) Dose-response curves of TWG-07-148 against Ba/F3 cell lines expressing BRAF^S365A^, CRAF^S259A^, and BRAF^V600E^. (**G**) Immunoblot analysis of Ba/F3 cells demonstrating inhibition of phospho-ERK by TWG-07-148 compared with trametinib. Note that trametinib is known to allow single-site phosphorylation of MEK on Ser222, and the single-phosphorylated MEK is recognized by most commercial anti-pMEK S218/222 antibodies(*29*).

**Figure 5.**
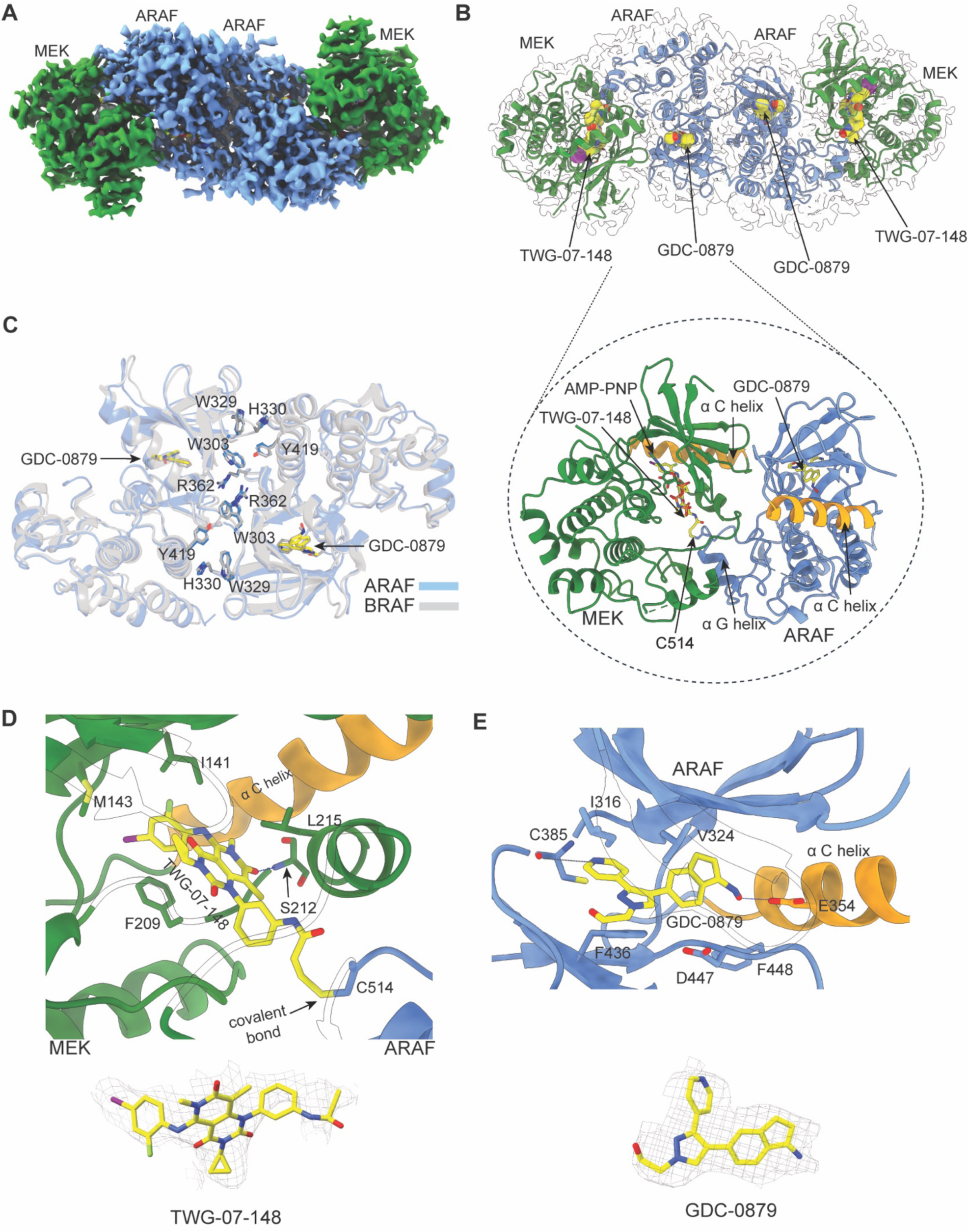
Cryo-EM structure of the ARAF-MEK1 heterotetramer bound to GDC-0879 and covalent MEK inhibitor TWG-07-148. (**A**) A 3.23 Å cryo-EM map of ARAF-MEK1 assembly reveals a heterotetrameric MEK1:ARAF:ARAF:MEK1 configuration. (**B**) Ribbon representation of the ARAF-MEK1 heterotetramer with the cryo-EM map shown as a transparent surface. GDC-0879 occupies the ARAF ATP binding pocket in both protomers, and TWG-07-148 occupies the MEK1-ARAF allosteric site. AMP-PNP occupies the ATP-site of both MEK1 protomers. The enlarged view highlights the ARAF-MEK interface. (**C**) Structural comparison of the ARAF back-to-back dimer interface in the present structure with that of active BRAF (from PDB entry 4MNF). Key dimer interface residues are identically conserved and are shown in blue for ARAF and gray for BRAF. Residue numbering corresponds to ARAF. (**D**) Detailed view of TWG-07-148 (yellow) in the allosteric pocket, where it forms a covalent bond with Cys514 in ARAF. Selected interacting residues are shown in stick form, and a hydrogen bond with the backbone amide of Ser212 is indicated by a dashed line. (**E**) Binding mode of GDC-0879 (yellow) within the ARAF active site. Key interacting amino acids are highlighted, and hydrogen bonds are shown as dashed lines. Cryo-EM density in the region TWG-07-148 and GDC-0879 is shown in the lower panels of d and e, respectively.

Due to the idiosyncratic increase in sensitivity of the ARAF^C514A^ to reversible inhibition with MEK inhibitors, this mutation was not useful for assessing the role of covalent bond formation in the increased potency of TWG-07-148 as compared with trametinib (Figure 4C and D, lower panels). However, in support of the importance of a covalent contribution to its potency, we note that replacement of the acrylamide in TWG-07-148 with the less reactive fluoroacrylamide warhead in TWG-07-178 resulted in a significant decrease in potency against ARAF^S214A^ (IC_50_ = 82.7 nM for TWG-07-178 versus 5.5 nM for TWG-07-148, Figure 3).

Although avutometinib derivatives GD-01-012 (acrylamide) and GD-01-014 (chloroacetamide warhead) efficiently labeled ARAF, they were less potent than the parent compound in the Ba/F3 cell assay, with IC_50_ values of approximately 450 nM (Figure 3). We did not observe efficient labeling with any of the cobimetinib or selumetinib derivatives, and these were not studied further. We therefore focused our further characterization on trametinib analog TWG-07-148, the only covalent compound that was demonstrably more potent than its reversible parent.

To determine whether covalent labeling of ARAF was MEK-dependent, i.e. whether engagement of the compound in the MEK allosteric pocket “delivered” the warhead to Cys514, we carried out a competition experiment with trametinib. In support of the intended mechanism of action, the presence of a 1:1 or 5:1 ratio of trametinib to TWG-07-148 proportionately blocked labeling of ARAF by TWG-07-148 when the ARAF-MEK complex was incubated with both compounds (Figure 4E).

We also profiled TWG-07-148 on our Ba/F3 cell models driven by BRAF^S365A^, CRAF^S259A^, and BRAF^V600E^ (Figure 4F, Supplementary Table 1). Measured potencies in the BRAF and CRAF cell lines closely paralleled those of trametinib; the enhanced potency of TWG-07-148 was unique to ARAF, which contains the target cysteine residue. Western blotting for phospho-MEK and phospho-ERK in these cell lines confirmed dose-dependent suppression of MAP kinase pathway signaling by TWG-07-148, comparable to that observed with trametinib (Figure 4G).

### Cryo-EM structure of an ARAF-MEK complex with TWG-07-148

To better understand the binding mode of TWG-07-148, we determined a cryogenic electron microscopy (cryo-EM) structure of the ARAF kinase domain in complex with MEK1, TWG-07-148, and the RAF inhibitor GDC-0879 (see Methods, Supplementary Figure 6, Supplementary Table 5). GDC-0879 is a type I RAF inhibitor that promotes the active conformation of the kinase and in turn drives dimerization of the kinase domain (*49*). Accordingly, with addition of GDC-0879 we observed a shift in elution volume of the ARAF-MEK1 complex on size exclusion chromatography (SEC) consistent with formation of a heterotetrameric complex corresponding to an ARAF_2_MEK1_2_ heterotetramer (MW ∼160 kDa, Supplementary Figure 6A). After incubation with TWG-07-148, this complex was used for cryo-EM imaging and single-particle reconstruction (Supplementary Figure 6B-E). The nominal resolution of the cryo-EM map was 3.2Å, but preferred orientation of the particles on the EM grid led to anisotropy in the reconstruction (Supplementary Figure 6F).

The resulting structure revealed a two-fold symmetric, heterotetrameric arrangement consisting of a back-to-back dimer of the ARAF kinase domain with MEK1 bound to each ARAF kinase (Figure 5A, B). To our knowledge, ARAF has not previously been resolved in its active state, but the dimer conformation and dimer interface is essentially the same as that observed in active BRAF and CRAF structures (Figure 5C). This is not surprising, considering that RAFs signal via formation of heterodimers as well as homodimers (*50–52*). The ARAF construct we used for structure determination incorporates the activating “SSDD” substitution of the NtA motif. However, as in available BRAF and CRAF structures, this region is not resolved in the ARAF cryo-EM density either.

The overall binding pose of TWG-07-148 is essentially the same as that of the non-covalent parent compound trametinib. The 4-bromo-2-fluorophenyl group binds in the back of the MEK allosteric pocket where it engages in hydrophobic interactions with Ile141 and Met143 (Figure 5D), and the oxygen atom of central pyrido-pyrimidine ring system is positioned to hydrogen bond with the main chain amide of Ser212 in MEK. Although the density is weak in the region of the acrylamide group of TWG-07-148, it is consistent with formation of the expected covalent bond with Cys514 in ARAF (Figure 5D). Contacts between ARAF and MEK in this region are closely similar to those seen in a recent structure of an inactive-state ARAF-MEK structure with NST-628 (*31*) and in other RAF-MEK complexes (*29*), indicating that covalent bond formation can be accomplished without significant main chain rearrangements in either ARAF or MEK.

The ARAF kinase domain exhibits a DFG-in, C-helix-in conformation, consistent with the type I binding mode of GDC-0879. GDC-0879 binds in the ARAF ATP-site in a manner analogous to that previously observed in BRAF (*53*). All residues that directly contact the compound are the same in ARAF and BRAF. The pyrazole core of GDC-0879 is sandwiched between Ile316 and Val324 in the P-loop and Phe436 in the C-lobe, and the pyridyl nitrogen hydrogen bonds with the backbone amide of Cys385 in the kinase hinge (Figure 5E). The ketoxime oxygen is positioned to hydrogen bond with Glu354 in the C-helix.

### TWG-07-148 inhibits proliferation of cancer cell lines driven by BRAF or KRAS mutations

We next evaluated the antiproliferative effects of TWG-07-148 on cancer cell lines harboring activating mutations in BRAF (HT29 and A375), KRAS (HT2122, A549, and HCT116) or EGFR (PC9). TWG-07-148 potently inhibited proliferation of both BRAF and KRAS mutant cancer cell lines, with IC_50_ values ranging from 6 nM for A375 (BRAF^V600E^) to 81 nM in A549 (KRAS^G12S^; Figure 6A,B). The compound was somewhat less potent against PC9 cells (168 nM), which harbor an EGFR exon 19 deletion. We profiled established MEK inhibitors trametinib, avutometinib, cobimetinib, and selumetinib in the same cell lines (Figure 6A,B). Unsurprisingly, the activity of TWG-07-148 most closely paralleled that of trametinib, with which it shares a common scaffold. In A549 cells, TWG-07-148 more comprehensively inhibited proliferation and was also modestly more potent as compared with trametinib. Similar to trametinib, TWG-07-148 inhibited phosphorylation of MEK and ERK in BRAF^V600E+^ melanoma cells lines A375 and HT29 (Figure 6C). Significant suppression of phospho-MEK and phospho-ERK was observed at concentrations as low as 0.01 µM, demonstrating potent on-target inhibition of signaling through the MAP kinase pathway.

**Figure 6.**
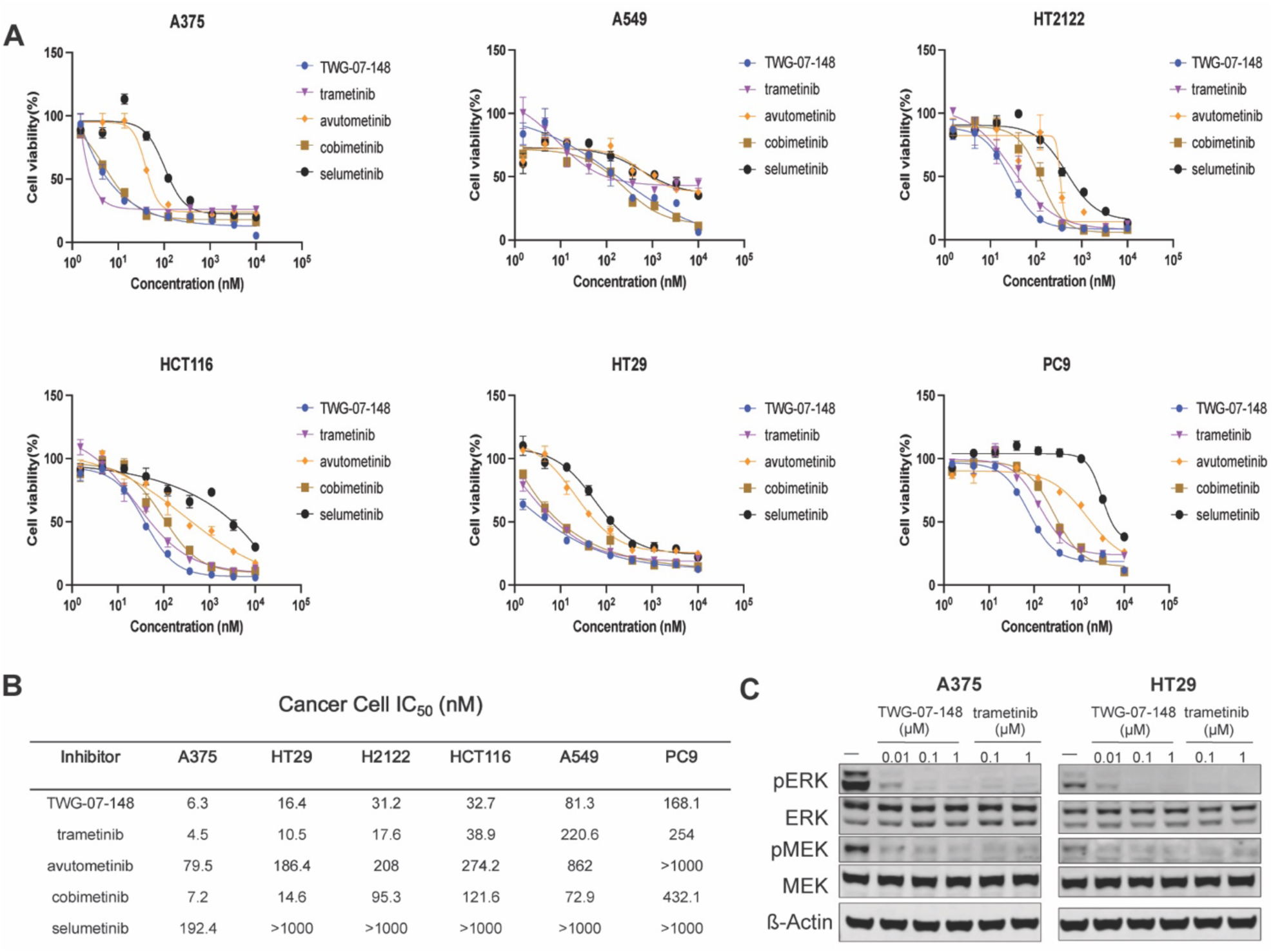
Profiling of TWG-07-148 and other allosteric MEK inhibitors in selected cancer cell lines. (**A**) Dose-response curves for TWG-07-148, trametinib, avutometinib, cobimetinib, and selumetinib against in A375, HT29, H2122, HCT116, A549, and PC9 cells. Cells were treated for 72 hours, and cell viability was assessed using the CellTiter-Glo assay. (**B**) Table of IC_50_ values derived from the dose-response curves in panel **A**. (**C**) Immunoblot analysis showing inhibition of phospho-MEK and phospho-ERK by TWG-07-148 and trametinib in A375 and HT29 cells.

Because MEK inhibitors have been shown to result in cell cycle arrest, we tested the effect of TWG-07-148 on cell cycle progression in both our RAF-mutant Ba/F3 cell lines and in BRAF^V600E+^ cancer cell lines. In ARAF^S214A^, BRAF^S365A^, and CRAF^S259A^ Ba/F3 cells, treatment with TWG-07-148 for 48 hours (at concentrations of 0.1 µM and 1 µM) significantly increased the fraction of cells in the G0/G1 phase, consistent with the expected effect of inhibiting the transition from G1 to the M phase (Supplementary Figure 7A, B). We observed a similar block when BRAF^V600E+^ A375 (melanoma) and HT29 (colorectal) cell lines were treated with TWG-07-148 or trametinib at the same concentrations (Supplementary Figure 7C, D).

## Discussion

Our profiling of seven allosteric MEK inhibitors across RAF isoforms reveals a previously unrecognized hierarchy of sensitivity, with all tested compounds showing greatest potency against CRAF-driven MEK activation and substantially reduced activity against ARAF-driven signaling. This isoform-dependent variation in inhibitor efficacy spanned 10- to 100-fold differences in potency and likely reflects an intrinsic property of RAF-MEK complexes, as it was observed in both cellular and biochemical assays. Through structure-guided mutagenesis, we identified mutations in the N-terminus of the RAF αG-helix that alter MEK inhibitor sensitivity. The dramatic sensitization to MEK inhibition in the ARAF C514A mutant, coupled with the resistance conferred by introducing cysteine or alanine at the corresponding position in CRAF, demonstrates that sequence variations in regions bordering the allosteric binding pocket can profoundly influence inhibitor efficacy. In addition, these findings further highlight the importance of the RAF partner as a determinant of sensitivity to MEK inhibition.

The mechanistic basis for the differences in RAF isoform sensitivity is clearly complex. Our mutagenesis experiments show that simply substituting the Gly-Cys-Arg sequence in the ARAF αG-helix with the Asn-Asn-Arg sequence found in CRAF does not confer comparable sensitivity to MEK inhibitors. Molecular dynamics simulations of the ARAF-MEK complex with ARAF C514A and C514N mutations show that both may have similar effects on the ARAF-MEK interface, despite their differing effects on inhibitor potency. The mechanism of action of MEK inhibitors is itself complex as it involves both stabilization of the MEK activation segment helix in a phosphorylation-resistant conformation and effects on the stability of the RAF-MEK interaction(*29*). Thus, differences across RAF isoforms in RAF-MEK affinity, in dynamic coupling between the RAF αG-helix and the MEK activation segment helix, as well as in direct contacts with the inhibitor can all contribute to observed drug sensitivity. These considerations also apply to the effects of the point mutations studied here.

We leveraged the presence of Cys514 in ARAF to develop TWG-07-148, a trametinib analog containing an acrylamide warhead that covalently targets this residue, which is unique to ARAF. This compound exhibits approximately 5-fold greater potency against ARAF-driven signaling compared to trametinib while maintaining comparable activity against BRAF and CRAF, providing proof-of-concept that the inherent ARAF-sparing tendency of current MEK inhibitors can be overcome through rational, covalent targeting of this residue.

The relative sparing of ARAF by current MEK inhibitors has important therapeutic implications, both positive and negative. The utility of anti-cancer therapeutics is always a balance between efficacy and tolerability, and a lack of potency on ARAF-MEK may contribute to the tolerability of MEK inhibitors because complete blockade of MAP kinase pathway signaling is not tolerated systemically. On the other hand, activating mutations in ARAF have been identified in Langerhans cell histiocytosis and in lung and other cancers(*37–39*), and existing MEK inhibitors may be less effective in these contexts as compared with BRAF or CRAF-driven cancers. Lack of potency against ARAF may also contribute to MEK inhibitor resistance. One widely observed mechanism of resistance to both RAF and MEK inhibitors is MAP kinase pathway reactivation due to relief of feedback inhibition (*54–56*). ERK signaling normally results in feedback inhibition of RAF activity, but RAF and MEK inhibitors can block this effect via RTK upregulation and consequent RAS and RAF activation promoting recovery of ERK activity. A lack of potency against ARAF may contribute to this phenomenon, as RAFs both homodimerize and heterodimerize and can cooperate in promoting resistance (*57*). The extent to which TWG-07-148 can mitigate resistance due to pathway reactivation is an important area for future study. Most current RAF inhibitors are also ARAF-sparing(*44*), which further emphasizes the potential value of more effective ARAF-MEK inhibitors.

Notwithstanding the potential confounding effects of RAF heterodimerization, ideally one would want to selectively block signaling from the specific RAF isoform activated in a given cancer. Our findings of varying potency of existing MEK inhibitors across RAF isoforms provide proof-of-principle for the feasibility of RAF-selective MEK inhibition, but we are not aware of deliberate efforts in the pharmaceutical industry to develop such an agent. Covalent targeting of Cys514 may offer an attractive approach to an ARAF-selective agent. While TWG-07-148 is not ARAF-selective due to its high-affinity reversible binding to BRAF-MEK and CRAF-MEK complexes, we speculate that a more selective ARAF-MEK inhibitor could be obtained by “de-tuning” its reversible affinity while maintaining the ability to form the covalent bond with ARAF.

Our study has several limitations. First, while our Ba/F3 cell models provide a controlled isogenic system for comparing MEK inhibitor potencies across RAF isoforms, these murine pro-B cells may not fully recapitulate the signaling context of human tumors driven by RAF or RAS mutations. Although we validated TWG-07-148 activity in human cancer cell lines harboring BRAF and KRAS mutations, we have not been able to identify an ARAF mutant or highly ARAF-dependent human cancer cell line that would allow us to more directly assess the advantage of its increased ARAF activity as compared with trametinib. Second, our biochemical assays employed 14-3-3-bound RAF kinase domain dimers that lack the regulatory regions of the full-length protein that influence RAF activity, localization, and complex formation in cells (*6*). Differences between cellular and biochemical potencies across RAF isoforms and for mutations such as ARAF C514A likely stem from factors beyond the immediate RAF-MEK interface mediated by these regions in the cellular milieu. Third, while the parent scaffold of TWG-07-148 is highly selective for MEK and our competition experiments support MEK-dependent delivery of the acrylamide warhead to Cys514, we have not assessed potential proteome-wide off-target reactivity or evaluated the compound’s selectivity across the kinome. Finally, we have only evaluated the activity of TWG-07-148 *in vitro* – we have not explored its pharmacokinetic properties or studied its efficacy *in vivo*.

In summary, our work establishes that RAF isoform identity is a critical determinant of MEK inhibitor potency, with current clinical agents exhibiting substantially reduced efficacy against ARAF-driven signaling. By identifying Cys514 as a unique structural feature of ARAF that can be exploited for covalent inhibition, we provide a rational strategy to overcome this limitation. While TWG-07-148 serves primarily as a proof-of-concept compound that will require further validation, it demonstrates that this covalent approach can meaningfully enhance inhibition of ARAF-MEK complexes. More broadly, our findings show that the view of MEK inhibitors as pathway-selective but otherwise non-discriminating agents is too simplistic; subtle differences in how these allosteric inhibitors engage different RAF-MEK complexes have functional consequences that could influence both therapeutic efficacy and resistance mechanisms. As our understanding of RAF isoform biology in cancer continues to evolve, the development of inhibitors tailored to specific RAF-driven contexts—whether through direct RAF kinase targeting or via RAF-selective MEK inhibition—represents a promising direction for improving outcomes in patients with MAP kinase pathway-driven malignancies.

## Materials and Methods

### Expression and purification of MEK1-RAF(A/B/C)-14-3-3 complexes for assay

The RAF proteins were expressed and purified as previously described (*44*). Briefly, BRAF(419-766) and CRAF(308-648) constructs containing their kinase domains and C-terminal 14-3-3 binding sites were expressed in a baculovirus mediated insect cell system. For ARAF(274-606), we used a previously described ARAF single-chain chimera in which 14-3-3 was genetically fused to the C-terminal end of the ARAF kinase domain (*44*). To express MEK1-RAF(A/B/C)-14-3-3 complexes, insect cells were coinfected with the corresponding baculoviruses. Cells were harvested 65–72 h postinfection and lysed by sonication in buffer A (50 mM Tris pH 8.0, 150 mM NaCl, 10 mM MgCl₂, 1 mM tris(2-carboxyethyl)phosphine (TCEP), 1 μM AMP-PNP), and Halt protease inhibitor cocktail (Thermo Fisher Scientific). Clarified lysates were loaded onto Ni-NTA agarose beads (Qiagen), washed with buffer A containing 30 mM imidazole, and eluted with buffer A containing 250–300 mM imidazole. ARAF and CRAF complexes were further purified on StrepTrap HP columns (Cytiva). Eluted complexes were concentrated to 1 ml (Amicon Ultra centrifugal filters 30 kDa MWCO, Millipore) and subjected to size-exclusion chromatography on Superose6 10/300 or Superdex200 Increase 10/300 column (Cytiva) for size-exclusion chromatography. SDS–PAGE confirmed co-elution of MEK1 with the respective RAF kinase domain and 14-3-3. The resulting MEK1:RAF:14-3-3 complexes exhibited kinase activity in TR-FRET assays. ERK2 expressions and purifications as substrates for assay were done as previously described (*44, 58*) using *E.coli* BL21(DE3) cells

### In vitro biochemical assay

Inhibition assays were performed using a modified cell-based HTRF phosphor-ERK kinase assay kit (Revvity). We purified ERK protein in-house and used it as our substrate. Inhibitors were dispensed into black 384-well plates using an HP300e dispenser and normalized to 1% final DMSO concentration per well. Kit assay buffer was supplemented with purified RAF at a final concentration of 2 nM for MEK1-BRAF-14-3-3 and MEK1-CRAF-14-3-3, and 10 nM for MEK1-ARAF-14-3-3𝜀, as well as purified ERK at a final concentration of 500 nM. 5μl of supplemented kinase buffer was dispensed into 384-well plates using a Multidrop combi dispenser (Thermo Fischer Scientific) and incubated with inhibitors at room temperature for 40 min before reactions were initiated by adding 5μl of 200 μM ATP dispensed . Plates were quenched after 2 hours at room temperature using the kit detection buffer supplemented with Anti-phospho ERK1/2 antibodies (Revvity) and were incubated overnight at room temperature. The FRET signal ratio was measured at 665 and 620 nm using a PHERAstar microplate reader (BMG Labtech) and processed using GraphPad Prism fit to a three-parameter dose-response model. Assays were performed in triplicate three independent times.

### Expression and purification of ARAF-MEK1 with GDC-0879 for cryo-EM

For structural studies, residues 272 to 579 of the human ARAF kinase domain with SSDD mutations were inserted into an insect cell expression vector with an N-terminal Maltose Binding Protein (MBP) tag and Tobacco Etch Virus (TEV)-cleavable His_6_ tag. Cells co-infected with MEK1 virus were harvested approximately 72 h postinfection via centrifugation and lysed in buffer B (pH 7.4, 20 mM HEPES, 300 mM NaCl, 5 mM MgCl_2_, 1mM TCEP, 1 μM AMP-PNP, 5% glycerol, 1μM GDC-0879) and protease inhibitor cocktail (Thermo Fisher Scientific) *via* sonication. Clarified lysates were loaded onto Ni-NTA agarose beads (Qiagen), washed with buffer B containing 30 mM imidazole, and eluted with buffer B containing 250 mM imidazole. The resulting fractions containing ARAF-MEK1 protein complex were concentrated and incubated with TEV protease (1:5 molar ratio) overnight to remove the MBP tag. ARAF-MEK1 protein complex was dialyzed against dialyzing buffer B.

The resulting sample was then subjected to another round of Ni-affinity purification (His-Trap, Cytiva) and the ARAF-MEK1 protein containing GDC-0879 inhibitor was eluted along a 100 ml gradient from 0 to 50% of buffer B containing 250mM Imidazole. Fractions containing ARAF-MEK1 were pooled and immediately injected onto Superdex 200 16/10 column for size exclusion chromatography. For plugning grids, the oligomeric ARAF-MEK1 (in complex with GDC-0879 inhibitor) peak fractions were concentrated to 0.8-1 mg/ml immediately after SEC and incubated overnight on ice at 4°C with covalent MEK inhibitor TWG-07-148 (protein to inhibitor 1:5 molar ratio), supplemented with additional 5mM MgCl2, 1μM AMP-PNP.

### Structure determination by cryo-EM

4μL of ARAF-MEK1 with both GDC-0879 and TWG-07-148 prepared as above was applied to a glow-discharged Quantifoil (Cu 1.2/1.3 400 mesh) grid, incubated for 10 seconds at 100% humidity and 10 °C, blotted for 5 seconds and plunge frozen in liquid ethane using a LEICA EM GP (Leica Microsystems). Images were acquired on a 300 kV Titan Krios microscope (Thermo Scientific) equipped with a Falcon4i direct electron detector and Selectris Energy Filter operated using EPU at Harvard Medical School cryo-Electron Microscopy Laboratory.

Images were collected with a total exposure time of 2 seconds, total dose of 51.09 e−/Å2, and a defocus range of -0.6 to -2.0 µm. Motion correction, CTF estimation, blob particle picking, and extraction were carried out in cryoSPARC (*59*). Particles were picked using an expected particle size of 80Å -180Å and a box size of 360 pixels. The super-resolution images were binned by 2 to the physical pixel size of 0.736Å during the extraction. A total of ∼12.4 million particles were selected by cryoSPARC and were subjected to extensive 2D classification until clear, distinguishable secondary structural features were visible. Following the 2D classifications, ∼1.3million particles were selected. Next, ab initio reconstructions were conducted with 4 classes while gradually increasing the maximum resolution from 12 to 8 Å. The resulting model comprised of ∼500k particles was used for heterogeneous refinement with C2 symmetry applied, while a non-protein-like-density was used as a decoy class. After a round of heterogeneous refinement coupled with one round of homogeneous refinement, ∼273k particles were selected for non-uniform refinement with low pass filter of 6 Å. From this set of particles (∼273k), another round of ab initio reconstructions was done with 4 classes to further refine this particle stack. Finally, 190k particles were selected for a final round of non-uniform refinement that ultimately yielded the final map. A final resolution of 3.23Å was determined using the gold-standard Fourier shell correlation (FSC)= 0.143 criterion using cryoSPARC(*59*) (Supplementary Figure 6F). The overall scheme for the data processing is summarized in Supplementary Figure 6D.

BRAF-MEK (PDB: 4MNE) crystal structure was used as the reference to build the ARAF-MEK1 model. The initial model was then fit into the cryo-EM density in UCSF ChimeraX 1.5(*60*). The model was refined against density map using Phenix 1.20 real space refinements with secondary structure restraints and no NCS constraints (*61*). The ARAF-MEK1 model was manually rebuilt and refined in the corresponding cryo-EM density map using Phenix 1.20 and WinCOOT 0.9.8 (*61, 62*). MolProbity was used to estimate the geometric restrains, clash score, and Ramachandran outliers (*63*).

### Generation of ARAF/BRAF/CRAF mutant Ba/F3 cells

Oncogenic kinase-transformed Ba/F3 cell lines were established as previously described(*64*). In brief, 3 μg of ARAF/BRAF/CRAF mutant expression vector was co-transfected with 3 μg of psPAX2 and 0.3 μg of pMD2.G (the packaging plasmids) into 70%-80% confluent 293T cells maintained in 3 mL of RPMI in a 60 mm dish by Lipofectamine 3000 (Thermo Fisher Scientific, cat# L3000001) mediated transfection. 48 hours post-transfection, the viral supernatant was harvested and filtered with a 0.45 μm membrane.

For infection of Ba/F3 cells, 2.5 mL of viral supernatant and 8 μg/mL polybrene was added to 2.5 million cells per well in six-well plates. Each plate was centrifuged at 2500 rpm for 90 minutes at 37 °C for the spin infection. 48 hours later, virus-containing medium was removed 48 hours later and successfully infected cells were selected with the addition of 1 μg/mL puromycin to the medium. After 3 days, cells were transferred into the fresh medium without interleukin-3 (IL-3) and puromycin. IL-3-independent transformed cells were maintained in RPMI medium 1640 supplemented with 10% FBS. Expression of ARAF/BRAF/CRAF were examined by Western blot.

### Cell culture

The ARAF/BRAF/CRAF mutant Ba/F3, H2122, PC9 and HT29 cells were cultured in RPMI media (Life technologies, Cat# 11875119) containing 10% fetal bovine serum (GeminiBio, Cat #100-106), 100 units/mL Penicillin and 100 μg/mL Streptomycin (gibco, cat #15140-22). A375 and A549 cells were cultured in DMEM media (Life technologies, Cat# 11995073) containing 10% fetal bovine serum (GeminiBio, Cat #100-106), 100 units/mL Penicillin and 100 μg/mL Streptomycin (gibco, cat #15140-22). HCT116 were cultured in McCoy’s 5A media (Life technologies, Cat# 16600082) containing 10% fetal bovine serum (GeminiBio, Cat #100-106), 100 units/mL Penicillin and 100 μg/mL Streptomycin (gibco, cat #15140-22). Parental Ba/F3 were cultured in RPMI media (Life technologies, Cat# 11875119) containing 10% fetal bovine serum (GeminiBio, Cat #100-106), 200ng of IL3 (Prospec, cat # CYT-371) and 100 units/mL Penicillin and 100 μg/mL Streptomycin (gibco, cat #15140-22).

All the cell lines were cultured at 37°C in 5% CO2-humidified air and tested for mycoplasma negative.

### Antiproliferation Assay

Cells were seeded at the density of 500 (adherent) or 1000 (suspension) cells per well in 384-well plates. For adherent cell lines, cells were cultured for 16 hours before compounds were added into the media. The compounds were dispensed into the cells by HP D300 digital drug printer and incubated for 72 h. Cell viability was determined by using CellTiter-Glo (Promega #G7571) according to the manufacturer’s instructions, measuring luminescence using a FLUOstar plate reader (BMG Labtech). Dose-response curves were generated using 3-parameter non-linear regression fits in GraphPad Prism(GraphPad Software). Data are presented as mean ± SEM (n = 3 or 4).

### Immunoblotting and Antibodies

Cells were lysed in RIPA buffer (150 mM NaCl, 1.0% IGEPAL® CA-630, 0.5% sodium deoxycholate, 0.1% SDS, 50 mM Tris, pH 8.0) (Sigma, cat# R0278) with protease and phosphatase inhibitor (Thermo Scientific, cat #1861282). The protein concentrations were measured by BCA analysis (Thermo Fisher Scientific, Cat # PI23225). Equal amounts of protein were resolved by 4-12% Tris-Base gels (Life Technologies) and then transferred to the Immuno-Blot PVDF membrane (BioRad, cat # 1620177). Proteins were probed with appropriate primary antibodies at 4 ℃ overnight and then with IRDye®800-labeled goat anti-rabbit IgG (LICOR Biosciences, cat # 926-32211), IRDye®800-labeled goat anti-mouse IgG (LICOR Biosciences, cat # 926-32210) or IRDye 680RD goat anti-Mouse IgG (LICOR Biosciences, Cat # 926-68070) secondary antibodies at room temperature for 1 hour. The membranes were imaged on an Odyssey CLx system.

Antibodies used in this study include anti-following proteins: p-MEK1/2 (Ser217/221) (Cell signaling Technology, #9154S, 1:1000), MEK1/2 (Cell signaling Technology, #8727S, 1:1000), ERK1/2 (Cell signaling Technology, #4696S, #1:0000), p-ERK1/2 (Cell Signaling Technology, #4370S, 1;1000), and β-Actin (Cell Signaling Technology, #3700, 1:1000).

### Cell cycle analysis

Cells were treated with DMSO, TWG-07-148 or Trametinib for 48 hr before harvesting and washing with cold PBS. The cells were then fixed with cold 70% ethanol (pre-chilled at -20°C) and incubated at -20°C overnight. On the day of flow cytometry analysis, cells were collected by centrifugation, wash cell pellets 3X with cold PBS, and stained in PI staining solution (25 μg/mL propidium iodide, 0.2 mg/mL RNase and 0.1% Triton X-100 in PBS) for 30 min at room temperature and then analyzed by LSRFortessa flow cytometer (BD Bioscience, NJ, USA). Results were analyzed by FlowJo 10.1 software (https://www.flowjo.com/solutions/flowjo)

### Chemical compounds

**TWG-07-148** *N*-(3-(3-cyclopropyl-5-((2-fluoro-4-iodophenyl)amino)-6,8-dimethyl-2,4,7-trioxo-3,4,6,7-tetrahydropyrido[4,3-d]pyrimidin-1(2H)-yl)phenyl)acrylamide

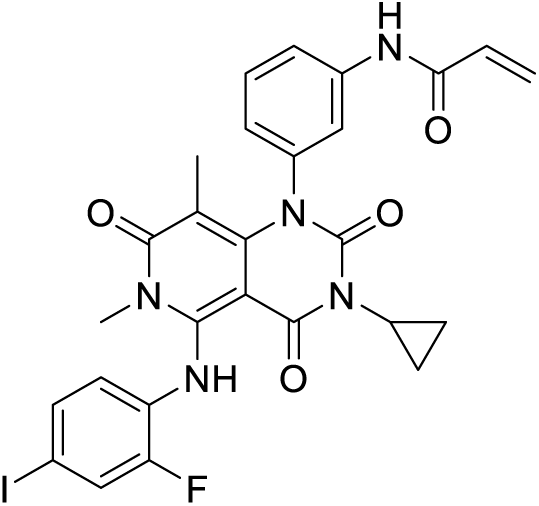

A mixture of 1-(3-aminophenyl)-3-cyclopropyl-5-((2-fluoro-4-iodophenyl)amino)-6,8-dimethylpyrido[4,3-d]pyrimidine-2,4,7(1H,3H,6H)-trione (30 mg, 0.056 mmol), HATU (43 mg, 0.112 mmol), acrylic acid (4.3 uL, 0.062 mmol), diisopropylethylamine (29 uL, 0.168 mmol), and DMF (0.5 mL) was stirred at RT for 1.5 hours. The reaction mixture was purified by reverse phase HPLC, 0-80% ACN/H_2_O (TFA modifier) to give the title compound (4.9 mg, 14 %). ^1^H NMR (500 MHz, DMSO-*d_6_*) δ ppm 11.08 (s, 1H) 10.32 (s, 1H) 7.79 (dd, 1H) 7.70 (m, 2H), 7.56 (dd, 1H) 7.42 (t, 1H) 7.09 (dd, 1H) 6.93 (t, 1H) 6.43 (dd, 1H) 6.27 (dd, 1H) 5.78 (d, 1H) 3.08 (s, 3H) 2.63 (m, 1H) 1.27 (s, 3H) 0.96 (m, 2H) 0.68 (m, 2H) 3 protons masked by solvent; MS *m/z*: 627.82 [M+1]^+^.

1-(3-Aminophenyl)-3-cyclopropyl-5-((2-fluoro-4-iodophenyl)amino)-6,8-dimethylpyrido[4,3-d]pyrimidine-2,4,7(1H,3H,6H)-trione was prepared according to the procedures in Abe et al, *ACS Med. Chem. Lett*. 2011, *2*, 320–324(*65*).

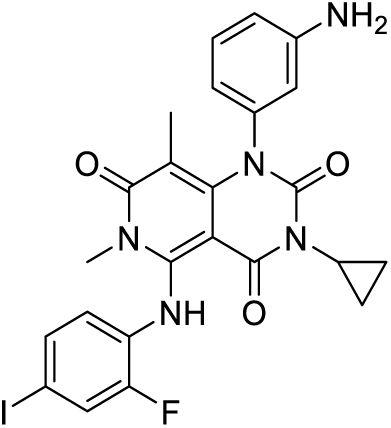

**GD-01-012** *N*-(2-fluoro-3-((4-methyl-2-oxo-7-(pyrimidin-2-yloxy)-2H-chromen-3-yl)methyl)phenyl) acrylamide

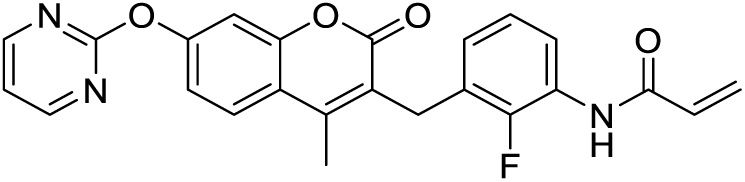

A solution of 3-(3-amino-2-fluorobenzyl)-4-methyl-7-(pyrimidin-2-yloxy)-2H-chromen-2-one (50 mg, 0.13 mmol), diisopropylethylamine (34.5 uL, 0.20 mmol), and DCM (2 mL) at 0°C was treated with acryloyl chloride (12.8 uL, 0.16 mmol). The reaction mixture was allowed to warm to room temperature and stirred for 15 minutes, then concentrated under reduced pressure. The residue was purified by reverse phase-HPLC, 0-80% ACN/H_2_O (TFA modifier) to give the title compound (46 mg, 81%). ^1^H NMR (500 MHz, DMSO-*d_6_*) δ: 9.92 (s, 1H) 8.69 (d, 2H) 7.92 (d, 1H) 7.87 (m, 1H) 7.37 (d, 1H) 7.34 (t, 1H) 7.27 (dd, 1H) 7.06 (t, 1H) 6.96 (m, 1H) 6.62 (dd, 1H) 6.28 (dd, 1H) 5.78 (dd, 1H) 4.03 (s, 2H), 3 protons masked by solvent peak; MS *m/z*: 432.11 [M+1]^+^.

3-(3-Amino-2-fluorobenzyl)-4-methyl-7-(pyrimidin-2-yloxy)-2H-chromen-2-one was prepared according to the procedures in Hyohdoh et al, *ACS Med. Chem. Lett*. 2013, *4,* 1059-1063(*66*).

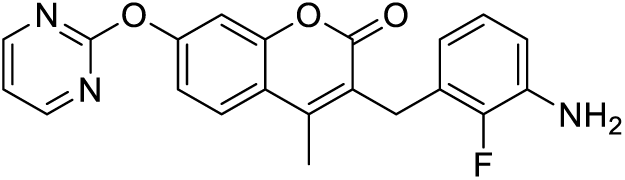

### Mass spectrometry analysis

Protein samples were acidified and desalted over C4 resin (ZipTip, Millipore). Samples were analyzed on an Orbitrap Eclipse mass spectrometer coupled to an Ultimate 3000 RSLCnano (Thermo Fisher Scientific). Proteins were separated over a 15-cm polystyrene divinylbenzene column (ES907, Thermo Fisher Scientific) with a 15-min gradient of 10-50% acetonitrile in 0.1% formic acid and electrosprayed (2 kV, 305°C) with an EasySpray ion source. Scans (600-2,000 m/z) were obtained in the orbitrap (15,000 resolution, profile). Deconvolution of the spectra and relative quantification of the peaks was done in the UniDec software(*67*). Percent labeling of proteins was calculated based on the relative ratio of labeled species to the parent species. Here, 3μM ARAF-MEK1 complex was treated with a five-fold excess concentration (1:5) of TWG-07-148 (15μM) or GD-01-012 (15μM).

### Molecular Dynamics Protocol

Initial structures were derived from the TWG-07-148 ARAF / MEK structure. Protein structures were prepared using the Protein Preparation Wizard in Schrödinger Suite (*68*). The TWG-07-148 covalent bond to ARAF C514 was broken. The terminal carbon of the acrylamide group was removed to convert TWG-07-148 into Trametinib. Mutate systems were prepared by muting residue 514 in Maestro (*64*). A local minimization of 0.5 nm around the Trametinib to remove possible steric clashes. Ligands were extracted from the system for parameterization. Ligand parameters were generated using the AMBER ff19SB force field (*69*). Small molecule structures were processed using AmberTools suite (*70*). AM1-BCC charges were calculated using Antechamber, with the GAFF2 force field parameters assigned via parmchk2. (*71*). Protein-ligand complexes were prepared by solvating the systems in a TIP3P water box with a padding of 1.2 nm. Counter-ions were added at a physiological ionic strength of 0.1 M. Energy minimization was performed using a convergence-based approach with a force tolerance of 1.0 kJ/mol/nm and an energy tolerance of 0.1 kJ/mol over 50 steps. The systems were equilibrated using a two-phase approach. A heating phase was conducted starting from 50 K and gradually increasing to 310 K over 2.5 ns, while then simultaneously reducing restraints on protein backbone atoms from 10.0 to 0.0 kJ/mol/nm² during the second 1.25 ns. Production simulations were performed in the NPT ensemble at 310 K and 1.0 bar, using the Langevin integrator with a friction coefficient of 1.0 ps⁻¹ and Monte Carlo barostat with a frequency of 25 steps. Hydrogen mass repartitioning (HMR) was used with a hydrogen mass of 4.0 Da, allowing for an extended timestep of 4.0 fs (*72*). All simulations were performed in triplicate with different initial velocities to ensure statistical significance, with each production run extending for 500 ns, yielding a total simulation time of 1.5 μs. Non-bonded interactions were truncated at 1.0 nm, and all covalent bonds involving hydrogen atoms were constrained using the SHAKE algorithm (*73, 74*). System stability during equilibration was confirmed by monitoring total energy and RMSD convergence. Simulations were analyzed via MDTRAJ (*75*). A separate post-processing protocol was implemented to quantify ligand–protein interaction energies from the production trajectories using OpenMM. For each saved frame, energy evaluations were performed, which were then averaged over the last 2500 frames to obtain mean and standard deviation values per system. All stages, from force-field assignment to simulation execution, were automated using custom Python workflows (available on GitHub). Scripts are available at https://github.com/dgazgalis/RAF-isoform-selectivity-of-MEK-inhibitors.

Software used in this study was curated by SBGrid (*76*).

## Acknowledgments

The authors thank members of the Eck lab and Jänne lab for helpful discussions and suggestions. We would also like to thank Milka Kostic for comments on the manuscript. Some of this work was performed at the Harvard Medical School cryo-EM facility, with the help from Dr. Richard Walsh. Part of the work was performed at Dana Farber Cancer Institute’s cryo-EM laboratory. Part of the work was performed at Dana Farber Cancer Institute’s mass spectrometry laboratory. Part of the work was also performed at Medicinal Chemistry core at Dana Farber Cancer Institute. We also thank SBGrid for the software curation.

## Author contributions

SC, JJ, and MJE designed the experiments; SC expressed and purified proteins, performed biochemical characterization of the inhibitors, analyzed the data, and determined the structure; JJ and FHG performed cell-based experiments, analyzed the data, performed immunoblotting, and cell sorting experiments; JVC performed initial protein expression trials and initial compound screening; DG ran molecular dynamics simulations; TWG, MT, and GD synthesized the covalent compounds; ET performed biochemical experiments; DMJ designed the constructs for protein expressions; BHH designed constructs and did initial protein expressional trials; TL assisted in the grid screening and structure determination; RJM and AS performed mass spectrometry-based experiments. DS, HJ, PAJ and MJE. contributed resources; HJ, PAJ, and MJE oversaw project. SC and MJE wrote the initial draft of the manuscript. All authors edited the manuscript.

## Data and materials availability

The data that support this study are available from the corresponding author upon request. All models and associated cryoEM maps have been deposited into the Electron Microscopy Data Bank (EMDBID: EMD-74071) and the Protein Data Bank (PDBID: 9ZDQ).

