## Supplemental Figures and Tables for "RAF isoform selectivity of MEK inhibitors and rational design of a covalent ARAF-MEK inhibitor"

Sayan Chakraborty et al.

**This PDF file includes:**

Figs. S1 to S7
Tables S1 to S5

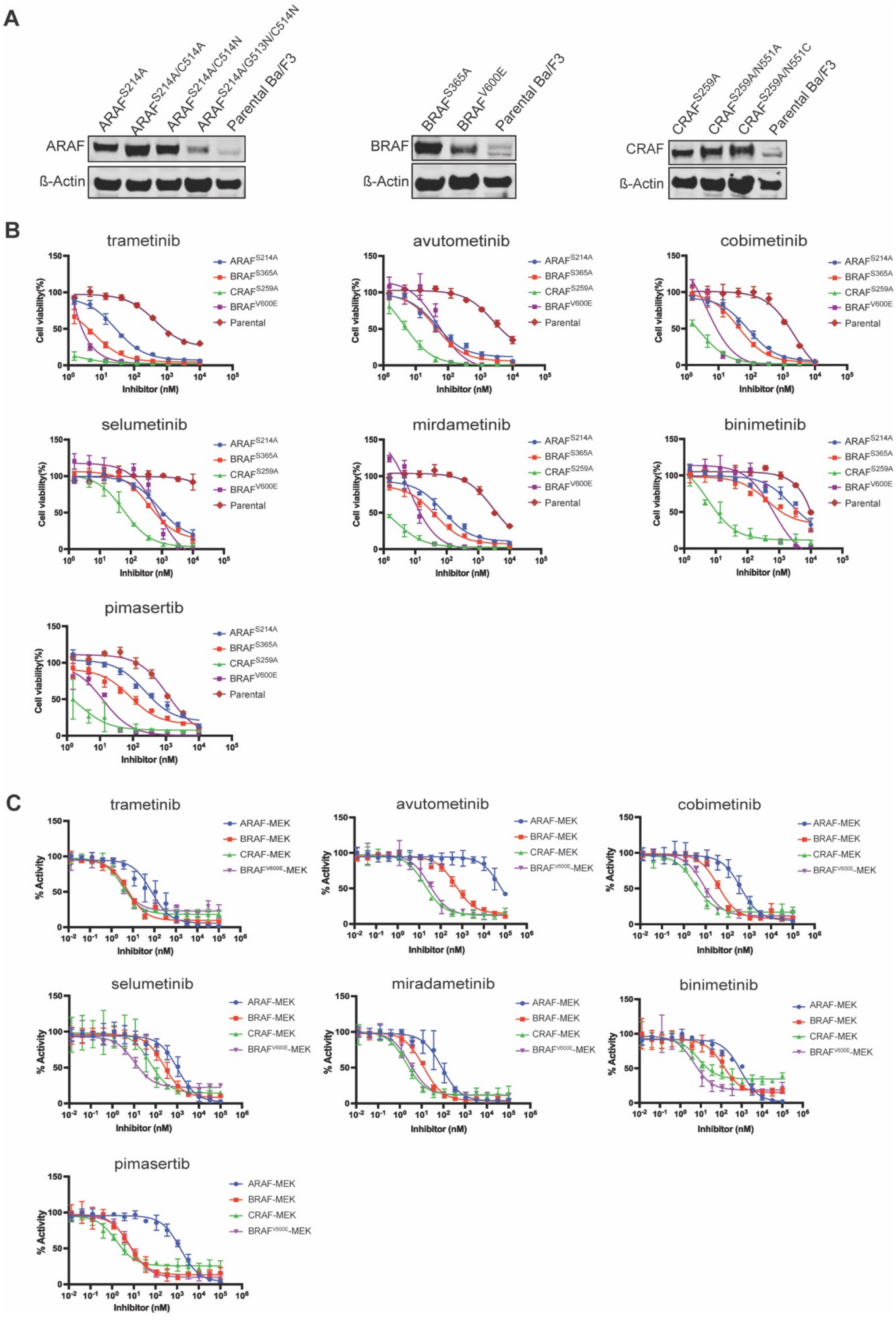
A
$ARAF^{S214A}$
$ARAF^{S214A/C514A}$
$ARAF^{S214A/C514N}$
$ARAF^{S214A/G513N/C514N}$
Parental Ba/F3
ARAF
ß-Actin
$BRAF^{S365A}$
$BRAF^{V600E}$
Parental Ba/F3
BRAF
ß-Actin
$CRAF^{S259A}$
$CRAF^{S259A/N551A}$
$CRAF^{S259A/N551C}$
Parental Ba/F3
CRAF
ß-Actin
B
trametinib
avutometinib
cobimetinib
selumetinib
mirdametinib
binimetinib
pimasertib
$ARAF^{S214A}$
$BRAF^{S365A}$
$CRAF^{S259A}$
$BRAF^{V600E}$
Parental
Cell viability(%)
Inhibitor (nM)
C
trametinib
avutometinib
cobimetinib
selumetinib
miradametinib
binimetinib
pimasertib
ARAF-MEK
BRAF-MEK
CRAF-MEK
$BRAF^{V600E}$-MEK
% Activity
Inhibitor (nM)

**Fig. S1. Ba/F3 cell expression of activated RAFs and dose response curves for allosteric MEK inhibitors across RAF isoforms.** (**A**) Immunoblot analysis of engineered Ba/F3 cells expressing constitutively active RAFs (ARAF$^{S214A}$, BRAF$^{S365A}$, CRAF$^{S259A}$, BRAF$^{V600E}$) and their corresponding mutant variants. (**B**) Dose-response curves for seven allosteric MEK inhibitors against engineered Ba/F3 cell lines expressing ARAF$^{S214A}$, ARAF$^{S214A/C514A}$, BRAF$^{S365A}$, V600E, and CRAF$^{S259A}$. (**C**) Dose response curves for seven allosteric MEK inhibitors in a biochemical cascade assay that measures phosphorylation of ERK2 by ARAF-MEK1, BRAF-MEK1, CRAF-MEK1, and BRAF$^{V600E}$-MEK1 complexes.

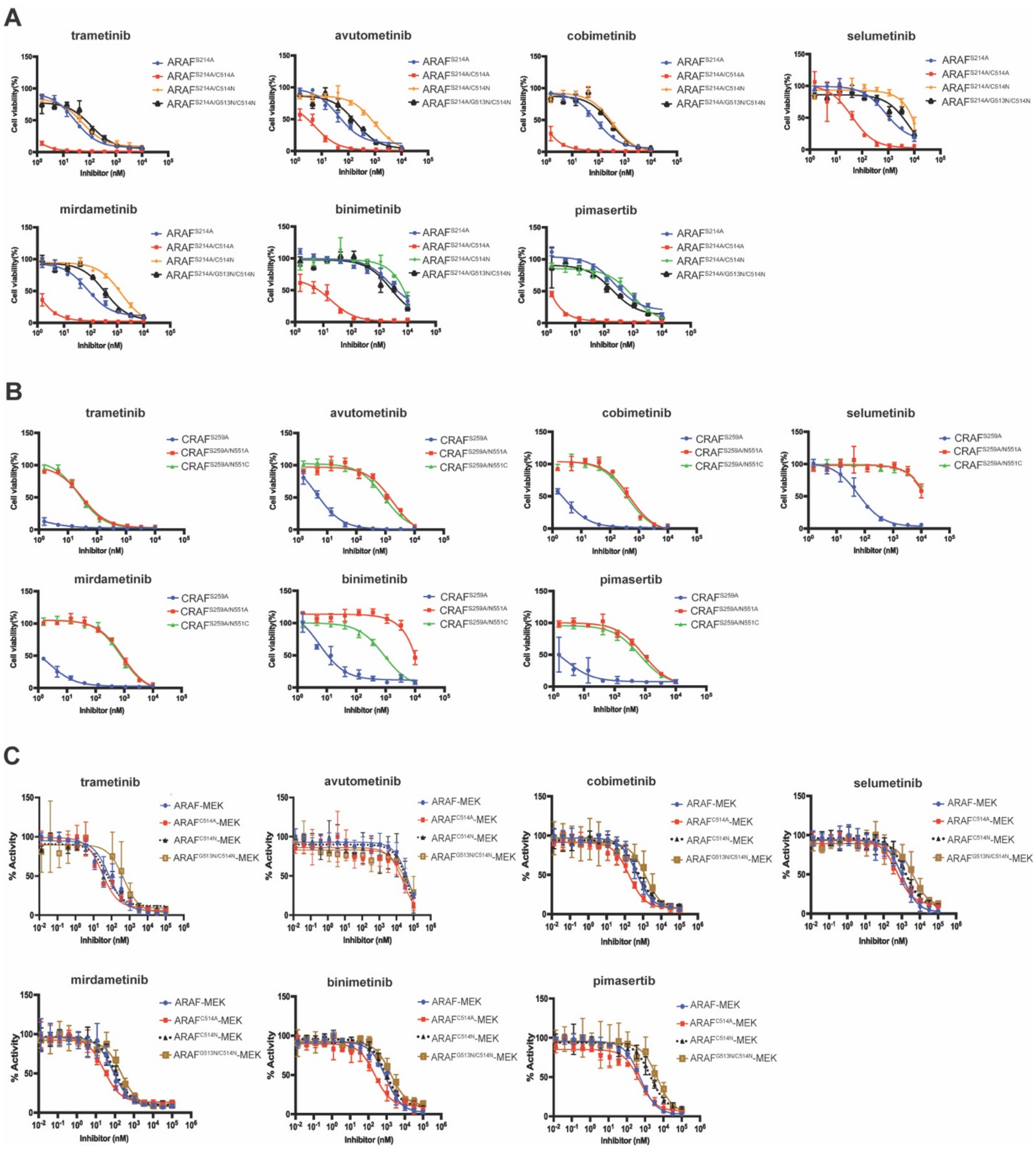


**Fig. S2. Dose response curves for allosteric MEK inhibitors against ARAF and CRAF αG-helix mutants.** (**A**) Cell viability assay assessing the potency of seven allosteric MEK inhibitors in engineered Ba/F3 cells expressing constitutively active ARAF$^{S214A}$ with and without mutations in the ARAF portion of the inhibitor binding pocket (αG-helix mutants C514A, C514N, and G513N/C514N). (**B**) Cell viability assay assessing the potency of allosteric MEK inhibitors in engineered Ba/F3 cells expressing constitutively active CRAF$^{S259A}$ with and without mutations in the CRAF portion of the inhibitor binding pocket (αG-helix mutants N553A and

N553C). (**C**) Dose response curves for allosteric MEK inhibitors in a biochemical cascade assay that measures phosphorylation of ERK2 by ARAF-MEK1, with or without mutations in the ARAF portion of the inhibitor binding pocket (αG-helix mutants C514A, C514N, and G513N/C514N).

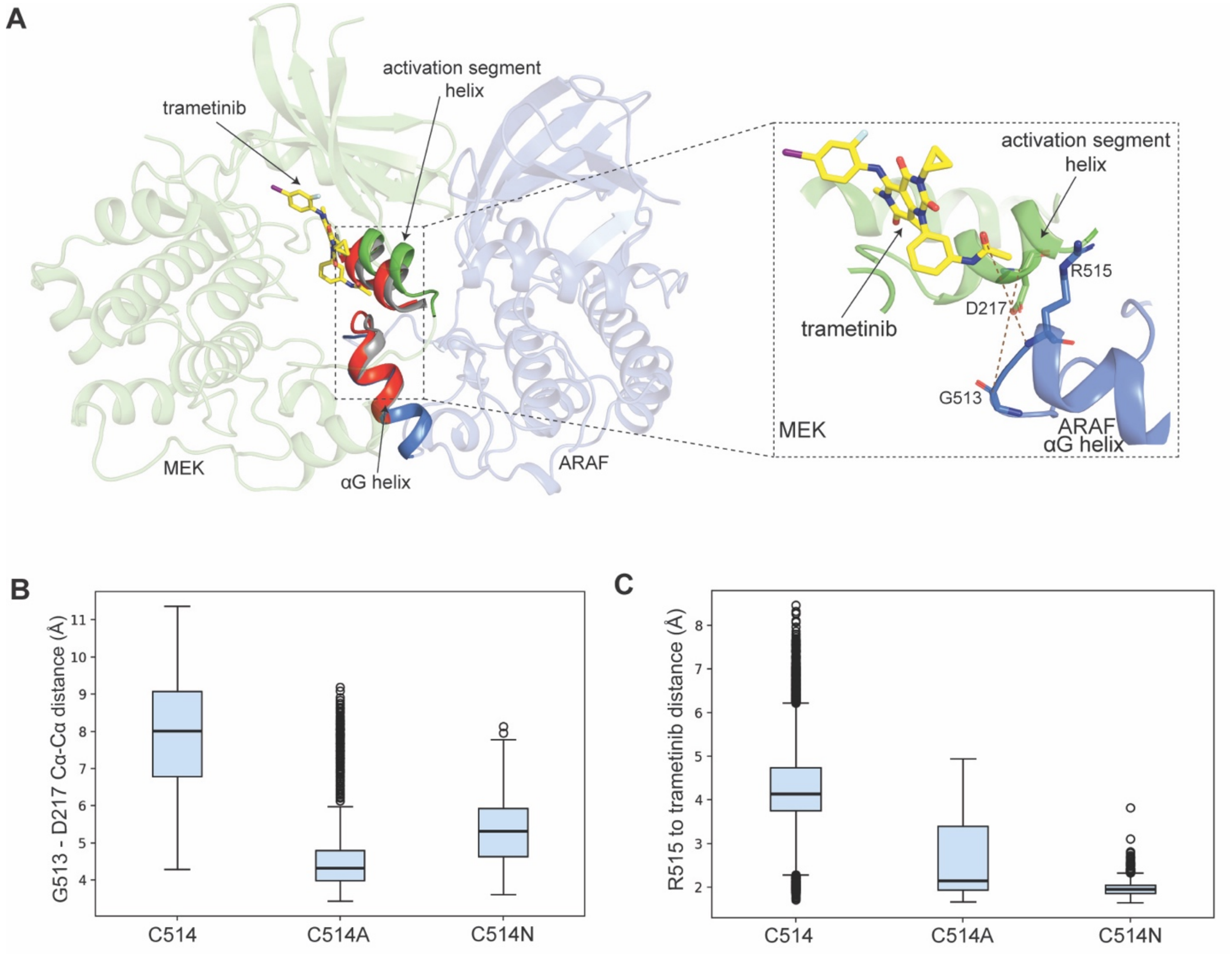


**D**

Calculated protein-ligand interaction energies

| System | Coulomb interaction (kcal/mol) | LJ interaction (kcal/mol) | Total Interaction (kcal/mol) |
|---|---|---|---|
| ARAF | -21.08 ± 5.22 | -71.90 ± 3.35 | -92.99 ± 6.07 |
| ARAF C514A | -27.91 ± 4.31 | -75.42 ± 3.19 | -103.33 ± 4.69 |
| ARAF C514N | -29.55 ± 4.03 | -71.58 ± 2.93 | -101.13 ± 4.66 |

**Fig. S3. Molecular dynamics analysis of the effect of ARAF αG-helix mutations on the ARAF-MEK protein complex and trametinib binding.** (**A**) Molecular dynamics simulations reveal subtle shifts in the positions of the activation segment helix of MEK1 and the N-terminal portion of the αG-helix of ARAF in the C514A and C514N mutants as compared with WT ARAF. Wild-type ARAF is shown in blue and MEK in green. Mutants C514A (red) and C514N (grey) are superposed on the wild-type ARAF-MEK complex. The inset illustrates the distance measurements plotted in panels b and c. (**B**) Box and whisker plots of the distance between ARAF G513 and MEK D217 (Cα-Cα) in wild-type versus C514A and C514N mutants reveal closer apposition of the MEK1 activation segment helix and the αG-helix of ARAF as seen in panel A. These distances were derived from MD simulation. (**C**) Box and whisker plots of the distance between the backbone amide NH of Arg515 in ARAF and the carbonyl oxygen of trametinib in wild-type versus C514A and C514N mutants indicate that this hydrogen bond is more stably formed in the ARAF mutants. (**D**) Calculated interaction energies between ARAF (wild-type and mutants) and trametinib obtained from the molecular dynamics simulations are shown in a tabulated form. Simulations were carried out for a total of 500 ns, analysis in **A**-**D** corresponds to mean position over final 250 ns of the simulation

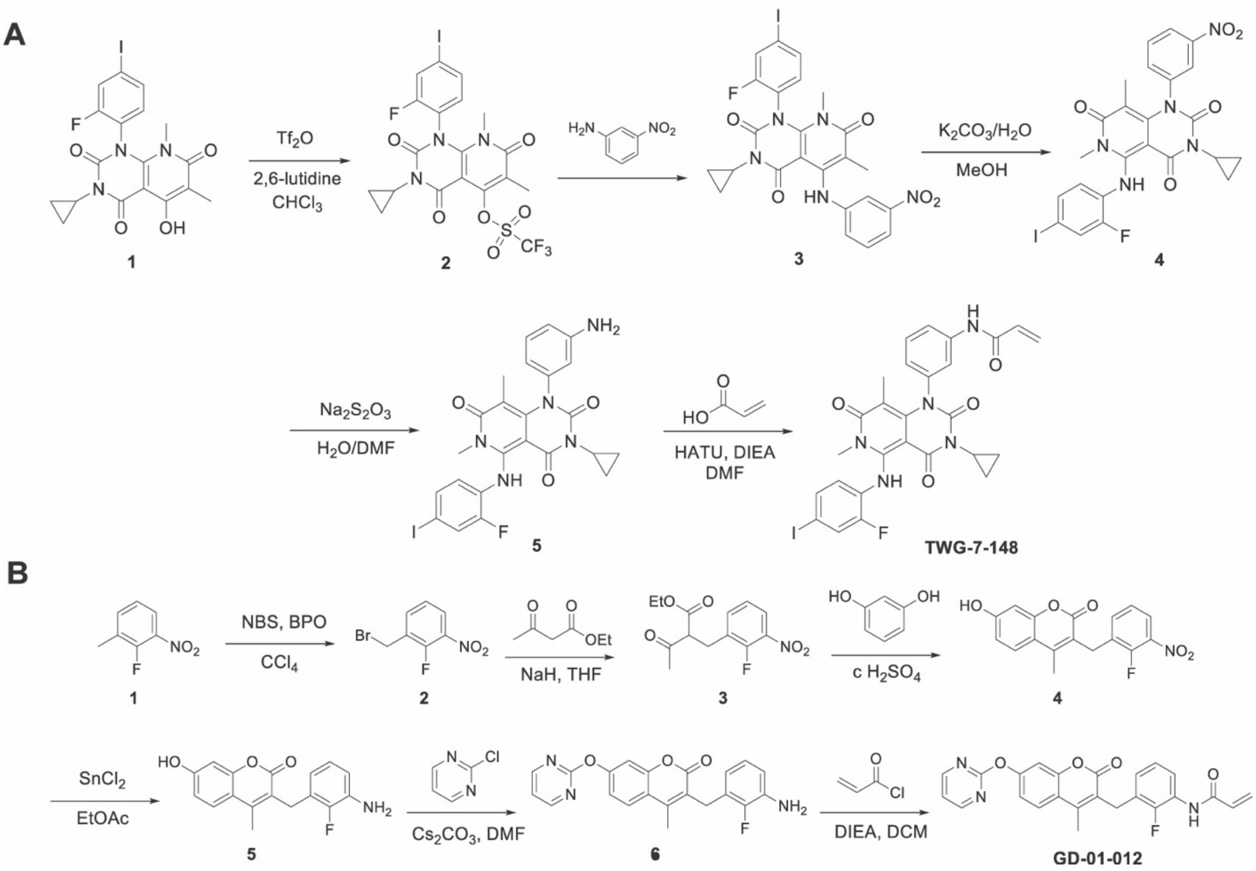


**Fig. S4. Synthetic scheme for TWG-07-148 and GD-01-012.** (**A**) Synthetic route for TWG-07-148. Compound **5** was prepared from 3-cyclopropyl-1-(2-fluoro-4-iodophenyl)-5-hydroxy-6,8-dimethylpyrido[2,3-d]pyrimidine-2,4,7(1H,3H,8H)-trione according to the procedures in Abe et al, ACS Med. Chem. Lett. 2011, 2, 320–324. (**B**) Synthetic route for GD-01-012. Compound **6** was prepared from 2-fluoro-3-nitrotoluene according to the procedures in Hyohdoh et al, ACS Med. Chem. Lett. 2013, 4, 1059-1063.

A

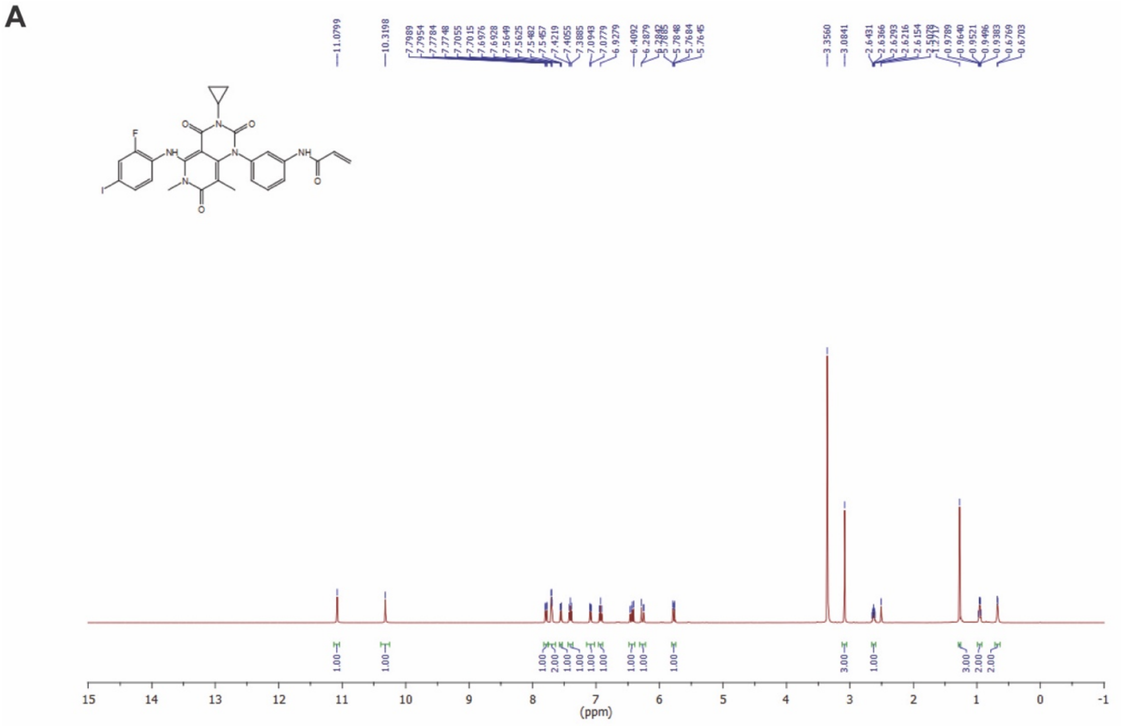


B

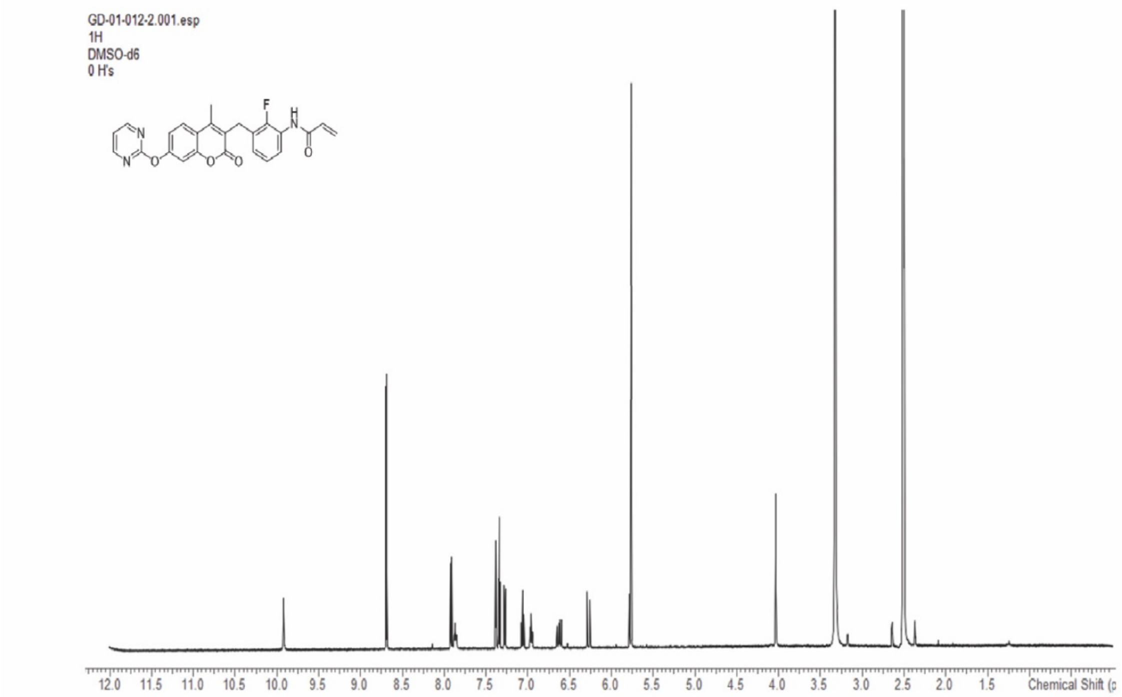


**Fig. S5. NMR spectra for TWG-07-148 and GD-01-012.** (**A** and **B**) $^1$H NMR spectra for purified TWG-07-148 and GD-01-012.

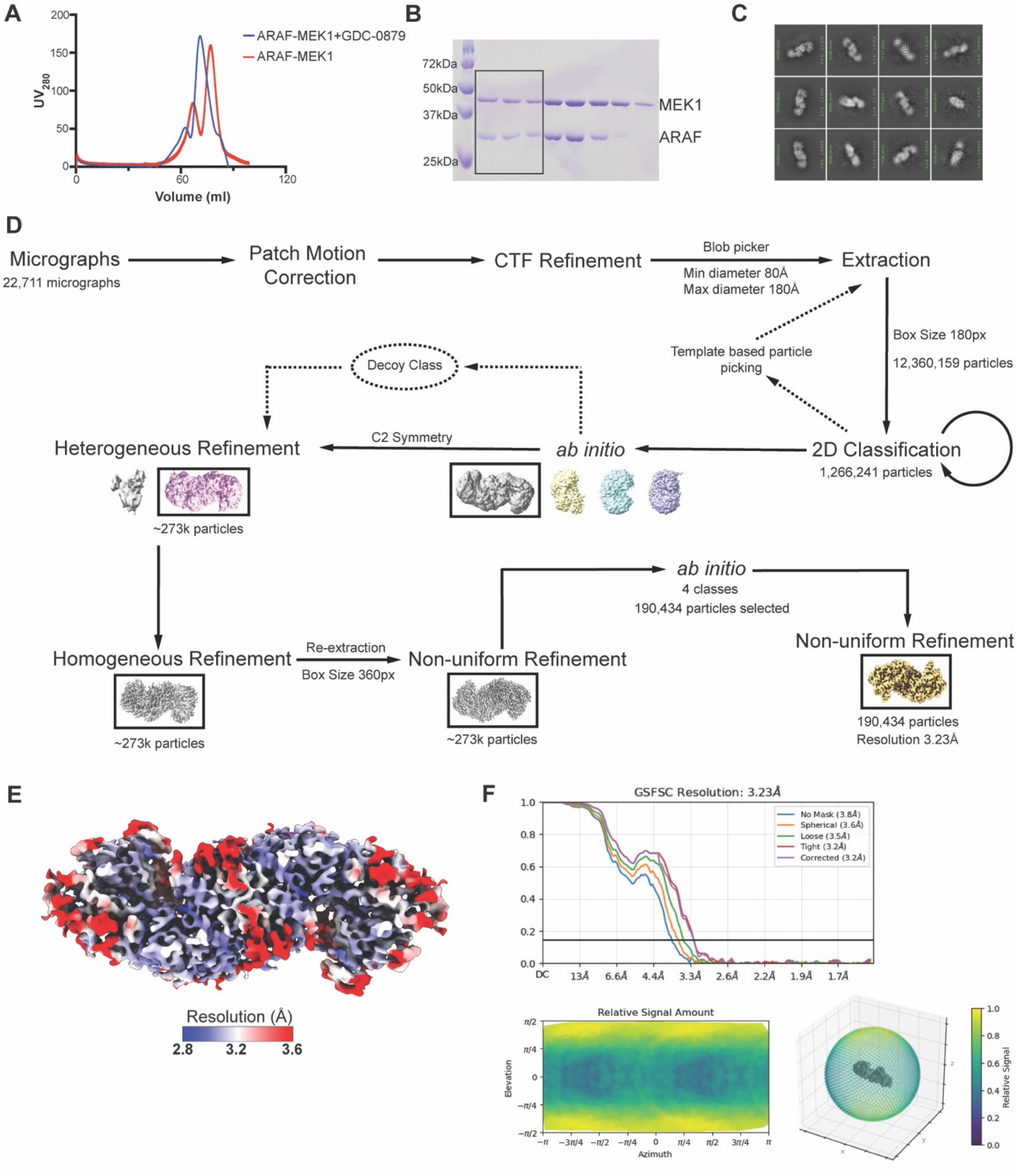


**Fig. S6. ARAF-MEK1 protein production and cryo-EM imaging scheme.** (**A**) Size exclusion chromatography (SEC) profile of purified ARAF-MEK1 with GDC-0879 (blue trace) or without added compound (red trace). Addition of type I inhibitor GDC-0879 shifts the elution profile to an earlier eluting peak, consistent with the expected conversion of an ARAF-MEK1 heterodimer to an $ARAF_2MEK1_2$ heterotetramer. Note that the ARAF construct corresponds to the kinase domain (residues 272 to 579, see *Methods* for further details). (**B**) SDS-PAGE analysis of SEC elution fractions obtained in the presence of GDC-0879. The red box indicates fractions that were pooled and used for cryo-EM imaging. (**C**) Representative 2D class averages of ARAF-MEK1 with GDC-0879 and TWG-07-148 obtained from CryoSPARC showing clear secondary structural features. (**D**) Schematic diagram showing the workflow for cryo-EM image processing. Image processing was done entirely in CryoSPARC. Classes with black boxes were selected for further processing. (**E**) Cryo-EM density map colored by local resolution as estimated in CryoSPARC. (**F**) Gold Standard Fourier shell correlation (GSFSC) curves showing the estimated resolution from CryoSPARC. The bottom panel illustrates the angular distribution of particle orientations and their contribution to the final cryo-EM reconstruction. The preferred orientations correspond to "side views" of the heterotetramer and resulted in noticeable anisotropy in the final map.

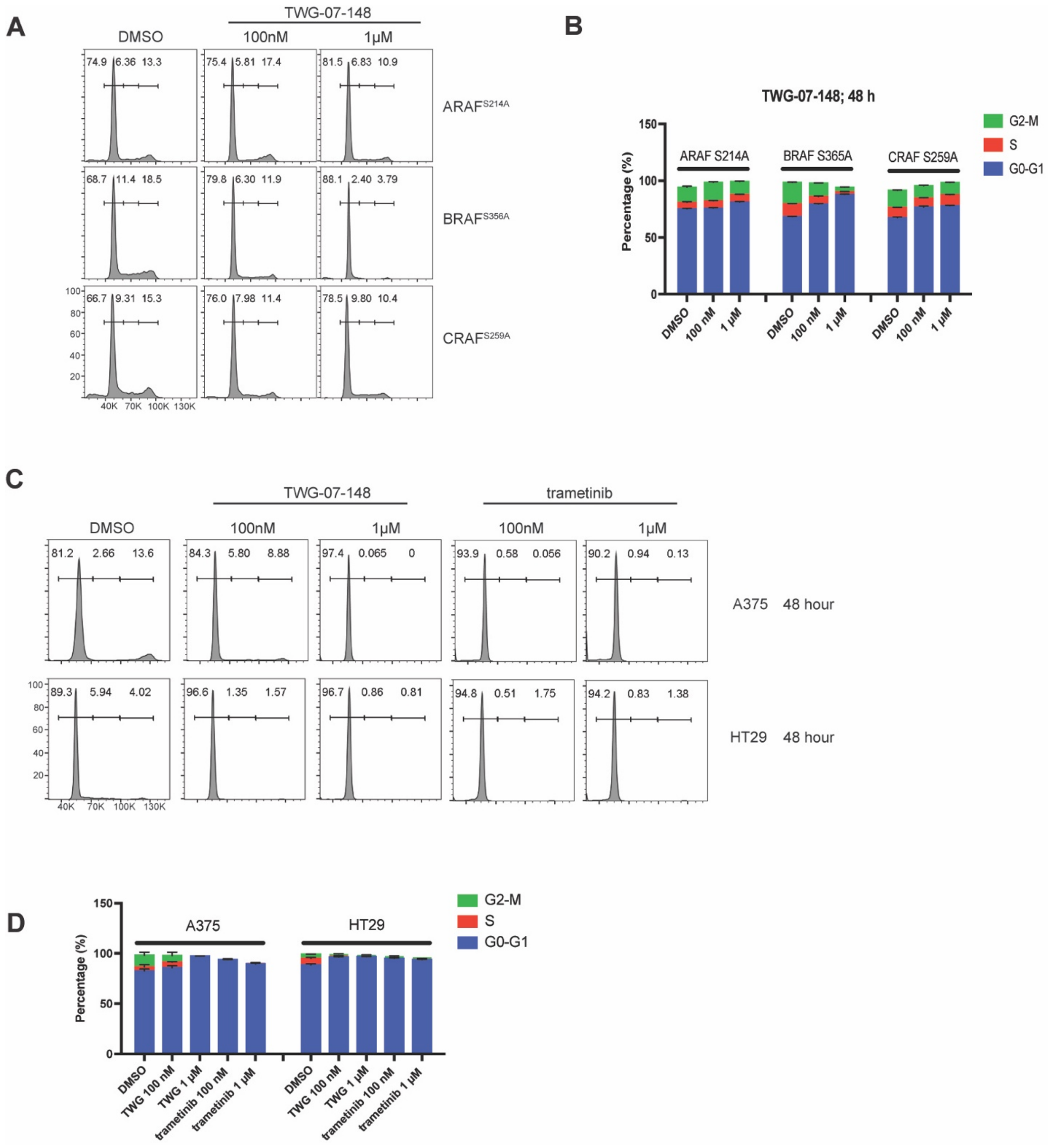


**Fig. S7. TWG-07-148 and trametinib induce cell cycle arrest.** (**A**) Flow cytometry cell cycle analyses of Ba/F3 cells transformed with ARAF$^{S214A}$, BRAF$^{S365A}$, or CRAF$^{S259A}$ and treated with TWG-07-148 (100 nM or 1 μM) or DMSO control. (**B**) Quantitation of fraction of cells in differing stages of the cell cycle ($G_0$-$G_1$, S, M) from panel **A**. (**C**) Flow cytometry cell cycle analyses of A375 and HT29 cells treated with TWG-07-148 (100 nM or 1 μM) or trametinib (100 nM or 1 μM). (D) Quantitation of fraction of cells in differing stages of the cell cycle ($G_0$-$G_1$, S, M) from panel c.

**Table S1. Ba/F3 cell $IC_{50}$ values for MEK inhibitors against constitutively active RAF isoforms.***

| Inhibitor | Ba/F3 cell $IC_{50}$ (nM) | | | | |
|---|---|---|---|---|---|
| | $ARAF^{S214A}$ | $BRAF^{S365A}$ | $CRAF^{S259A}$ | $BRAF^{V600E}$ | Parental |
| trametinib | 28.9 | 3.8 | <1 | 3.5 | >1000 |
| avutometinib | 69 | 48.5 | 5.5 | 72.6 | >1000 |
| cobimetinib | 78 | 46.7 | 2.1 | 11 | >1000 |
| mirdametinib | 94.7 | 34.7 | 1 | 12.1 | >1000 |
| selumetinib | >1000 | 734.2 | 69 | 856.5 | >1000 |
| binimetinib | >1000 | 117.2 | 17.2 | 605.2 | >1000 |
| pimasertib | 127.8 | 70.8 | <1 | 15.6 | >1000 |
| TWG-07-148 | 5.5 | 3.3 | <1 | 4 | 709 |

*$IC_{50}$ values correspond to dose response data plotted in Supplementary Fig. S1B and Fig. 1A in the main text.

**Table S2. Biochemical $IC_{50}$ values (nM) for MEK inhibitors against purified RAF-MEK complexes.***

| Inhibitor | ARAF-MEK | BRAF-MEK | CRAF-MEK | $BRAF^{V600E}$-MEK |
|---|---|---|---|---|
| trametinib | 72.9 | 7.5 | 5.4 | 5.1 |
| avutometinib | >1000 | 491 | 17 | 64 |
| cobimetinib | 324.3 | 47.6 | 2.8 | 3.3 |
| mirdametinib | 49.6 | 10.6 | 1.8 | 1.5 |
| selumetinib | >1000 | 209 | 41.7 | 17.2 |
| binimetinib | >1000 | 119.8 | 13 | 9.1 |
| pimasertib | 855 | 41.1 | 3.1 | 3 |

*$IC_{50}$ values correspond to dose response data plotted in Supplementary Fig. S1C and Fig. 1C in the main text.

**Table S3. Ba/F3 cell $IC_{50}$ values for MEK inhibitors against ARAF and CRAF variants.***

| | $ARAF^{S214A}$ $IC_{50}$ (nM) | | | | $CRAF^{S259A}$ $IC_{50}$ (nM) | | |
|---|---|---|---|---|---|---|---|
| **Inhibitor** | **C514** | **C514A** | **C514N** | **G513N/C514N** | **N553** | **N553A** | **N553C** |
| trametinib | 28.9 | <1 | 108.6 | 79.8 | <1 | 44.9 | 43.6 |
| avutometinib | 69 | 8.3 | 765 | 261.8 | 5.5 | >1000 | >1000 |
| cobimetinib | 78 | 4 | 333 | 259.6 | 2.1 | 499.5 | 503.2 |
| mirdametinib | 94.7 | 2.7 | 960 | 514.2 | 1 | 931 | 908.1 |
| selumetinib | >1000 | 2.8 | >1000 | >1000 | 69 | >1000 | >1000 |
| binimetinib | >1000 | 5.2 | >1000 | >1000 | 17.2 | >1000 | >1000 |
| pimasertib | 127.8 | 1.3 | 690.6 | 473.3 | <1 | >1000 | 990.2 |
| TWG-07-148 | 5.5 | <1 | 89.7 | 55.07 | <1 | 145.2 | 137.8 |

*$IC_{50}$ values correspond to dose response data plotted in Supplementary Fig. S2A, B and Fig. 2C, D in the main text.

**Table S4. Biochemical $IC_{50}$ values for MEK inhibitors against purified ARAF variants.***

| Inhibitor | ARAF-MEK $IC_{50}$ (nM) | | | |
|---|---|---|---|---|
| | C514 | C514A | C514N | G513N/C514N |
| trametinib | 72.9 | 25.9 | 34.9 | 18 |
| avutometinib | >1000 | >1000 | >1000 | >1000 |
| cobimetinib | 324.3 | 181.1 | 337.3 | 765 |
| mirdametinib | 49.6 | 29.6 | 53.5 | 130.6 |
| selumetinib | >1000 | 539.2 | >1000 | >1000 |
| binimetinib | >1000 | 296.5 | 651.4 | >1000 |
| pimasertib | 855 | 523.3 | >1000 | >1000 |

*$IC_{50}$ values correspond to dose response data plotted in Supplementary Fig. S2C.

**Table S5. Cryo-EM data collection, refinement, and validation statistics**

| **Data collection and processing** | **PDB : 9ZDQ**<br>**EMDB : 74071** |
|---|---|
| Magnification | 165,000x |
| Voltage (kV) | 300 |
| Electron exposure (e–/Å2) | 51.09 |
| Defocus range (μm) | -0.6 to -2.0 |
| Pixel size (Å) | 0.736 |
| Symmetry imposed | C2 |
| Initial particle images (no.) | 12,360,159 |
| Final particle images (no.) | 190,434 |
| Map resolution (Å) | 3.23 |
| FSC threshold | 0.143 |
| Map resolution range (Å) | 3/3.4/3.8 |
| **Refinement** | |
| Initial model used (PDB code) | 4MNE |
| Model resolution (Å) | 3.4 |
| FSC threshold | 0.143 |
| Model resolution range (Å) | 3.1/3.2/3.6 |
| Map sharpening B factor (Å2) | 119.1 |
| **Model composition** | |
| Non-hydrogen atoms | 8756 |
| Protein residues | 1082 |
| Ligands | T14:2/29L:2/ANP:2 |
| **B factors (Å2)** | |
| Protein | 21.45/166.00/83.83 |
| Ligand | 48.20/139.73/71.35 |
| **R.m.s. deviations** | |
| Bond lengths (Å) | 0.003 |
| Bond angles (°) | 0.548 |
| **Validation** | |
| MolProbity score | 2.32 |
| Clashscore | 7.27 |
| Poor rotamers (%) | 5.30 |
| **Ramachandran plot** | |
| Favored (%) | 94.54 |
| Allowed (%) | 5.46 |
| Outlier | 0.00 |
